# A small molecule VHL molecular glue degrader for cysteine dioxygenase 1

**DOI:** 10.1101/2024.01.25.576086

**Authors:** Antonin Tutter, Dennis Buckley, Andrei A. Golosov, Xiaolei Ma, Wei Shu, Daniel J. J. McKay, Veronique Darsigny, Dustin Dovala, Rohan Beckwith, Jonathan Solomon, Pasupuleti Rao, Lei Xu, Aleem Fazal, Andreas Lingel, Charles Wartchow, Jennifer S. Cobb, Amanda Hachey, Jennifer Tullai, Gregory A. Michaud

## Abstract

The Von Hippel-Lindau Tumor Suppressor gene product (pVHL) is an E3 ligase substrate receptor that binds proline-hydroxylated HIF1-α, leading to its ubiquitin-dependent degradation. By using protein arrays, we identified a small molecule that binds the HIF1-α binding pocket on pVHL and functions as a molecular glue degrader of the neosubstrate cysteine dioxygenase (CDO1) by recruiting it into the VHL-cullin-ring E3 ligase complex and leading to its selective degradation. The CDO1 binding region involved in VHL recruitment was characterized through a combination of mutagenesis and protein-protein docking coupled with molecular dynamics-based solvation analysis. The X-ray structure of the ternary complexes of VHL, CDO1, and degrader molecules confirms the binding region prediction and provides atomic insights into key molecular glue interactions.

---

The Von Hippel-Lindau tumor suppressor (pVHL) is the substrate recognition component of a cullin-ring ligase complex that regulates the stability of HIF1-α^1^. HIF1-α, as a heterodimer with HIF1-β, forms the active HIF-1 transcription factor which controls the expression of many genes important for adaptation under conditions of hypoxia. During normoxic conditions, specific HIF-1 prolyl hydroxylases bring about proline hydroxylation of HIF1-α which leads to high affinity binding to pVHL. Subsequently, the pVHL-cullin ring ligase complex brings about ubiquitination of HIF1-α leading to proteasomal degradation. Since the first reports of peptidomimetic VHL ligands based on the hydroxyproline motif/moiety in HIF^2–4^, these ligands have been further optimized to compound **4**^5^ (**Table 1**). Since the report of compound **4**, this core structure has served as the basis for a multitude of bifunctional degraders, or PROTACs^6–8^ such as compound **1**/MZ1^9^ (**Table 1**), which links the VHL ligand to JQ1 to yield a potent degrader of BET bromodomains. More recently, it was demonstrated that linking the KRAS^G12C^ covalent inhibitor MRTX849 to a VHL binding ligand leads to degradation of endogenous KRAS^G12C^ in cancer cell lines^10,11^. While these PROTACs have enjoyed many recent successes as *in vitro* and *in vivo* chemical probes, their large size and properties can present challenges for their development as drugs. In contrast to PROTACs, molecular glues more often resemble conventional drug-like small molecules and thereby can overcome several shortcomings of PROTACs. Smaller molecules are more likely to have favorable membrane permeability and cellular uptake. Their smaller size also makes the chemistry more synthetically tractable. Importantly, molecular glues demonstrate inherently high binding cooperativity with their substrates compared with PROTACs, thereby avoiding a central affinity-kinetics challenge intrinsic to PROTACs, namely the so-called hook effect wherein high concentrations of a PROTAC inhibit ternary complex formation^12^. Yet, *bona fide* molecular glue degraders have remained elusive, and to date no high cooperativity small molecule glue degraders for VHL have been reported. In this report, we describe the discovery of a small molecule that facilitates the assembly of a VHL-small molecule-CDO1 ternary complex that brings about VHL-dependent degradation of endogenous CDO1. The X-ray structure of the ternary complexes reveal key interactions that drive the recruitment of CDO1 to the novel protein interaction surface created on VHL by the degrader molecules.

**Table 1.**
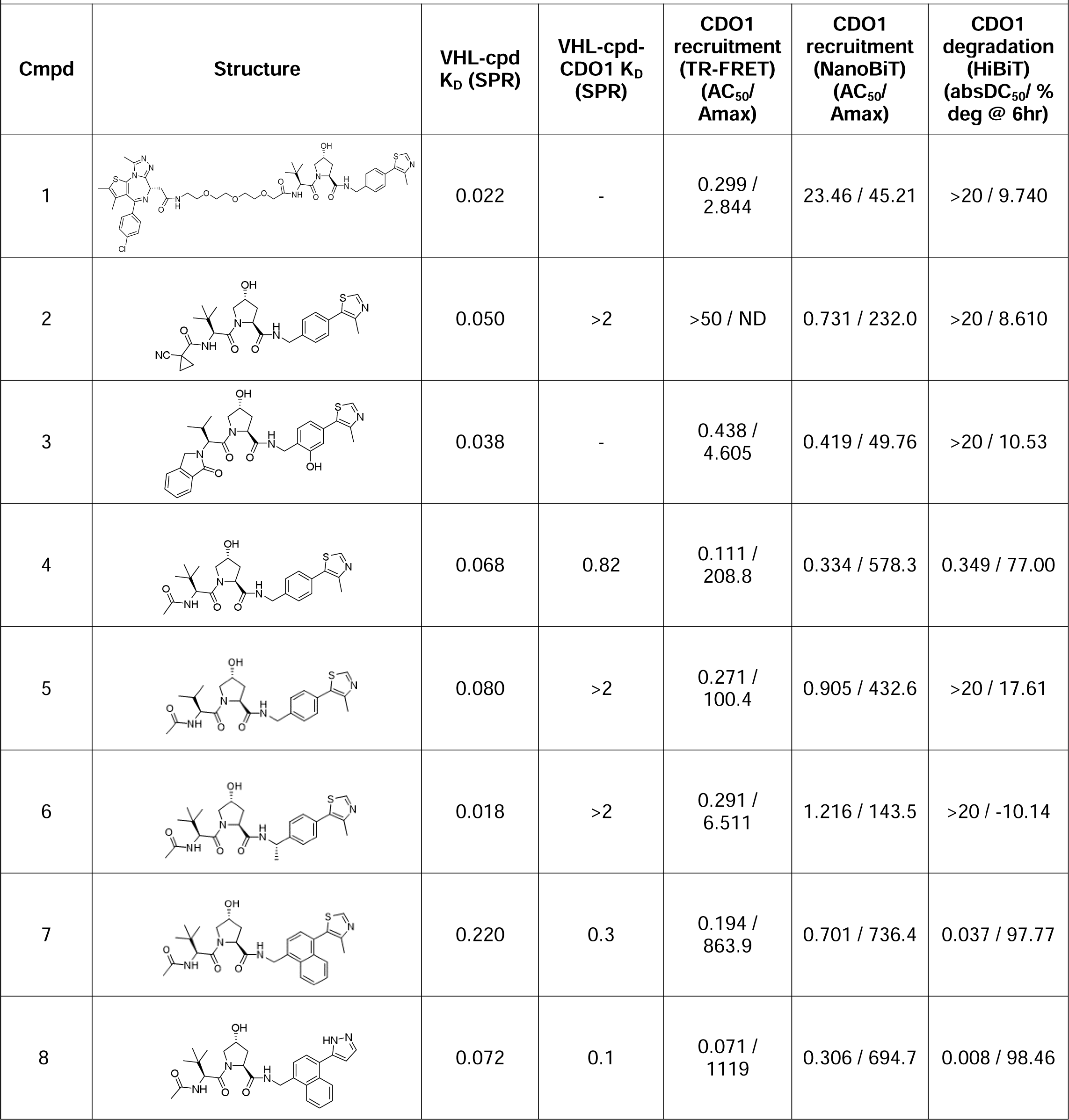
Summary of compounds used throughout this study. For each compound, the binding constant (K_D_) for its interaction with VHL (binary) and with the VHL-cpd-CDO1 (ternary) complex is indicated, in µM. CDO1 recruitment (AC_50_/A_max_) to VHL, by TR-FRET and NanoBiT, are shown. AC_50_ units are µM. A_max_ units are % activity above control. For compound-mediated degradation of CDO1 measured by HiBiT detection, degradation is indicated by the compound degradation absDC_50_, and by the maximum percent reduction in CDO1 protein level achieved. absDC_50_ units are µM. ND = not determinable.

| Cmpd | Structure | VHL-cpd<br>K<sub>D</sub> (SPR) | VHL-cpd-CDO1 K<sub>D</sub><br>(SPR) | CDO1<br>recruitment<br>(TR-FRET)<br>(AC<sub>50</sub>/<br>Amax) | CDO1<br>recruitment<br>(NanoBiT)<br>(AC<sub>50</sub>/<br>Amax) | CDO1<br>degradation<br>(HiBiT)<br>(absDC<sub>50</sub>/ %<br>deg @ 6hr) |
| --- | --- | --- | --- | --- | --- | --- |
| 1 |  | 0.022 | - | 0.299 /<br>2.844 | 23.46 / 45.21 | >20 / 9.740 |
| 2 |  | 0.050 | >2 | >50 / ND | 0.731 / 232.0 | >20 / 8.610 |
| 3 |  | 0.038 | - | 0.438 /<br>4.605 | 0.419 / 49.76 | >20 / 10.53 |
| 4 |  | 0.068 | 0.82 | 0.111 /<br>208.8 | 0.334 / 578.3 | 0.349 / 77.00 |
| 5 |  | 0.080 | >2 | 0.271 /<br>100.4 | 0.905 / 432.6 | >20 / 17.61 |
| 6 |  | 0.018 | >2 | 0.291 /<br>6.511 | 1.216 / 143.5 | >20 / -10.14 |
| 7 |  | 0.220 | 0.3 | 0.194 /<br>863.9 | 0.701 / 736.4 | 0.037 / 97.77 |
| 8 |  | 0.072 | 0.1 | 0.071 /<br>1119 | 0.306 / 694.7 | 0.008 / 98.46 |

## Results

### Protein microarray screening identifies a VHL molecular glue for CDO1

To identify novel substrates that get recruited to VHL in the presence of VHL small molecule ligands, we leveraged protein microarrays. First, we developed a protocol to detect VHL-small molecule induced PPIs using the PROTAC molecule compound **1**/MZ1 (**Table 1**) and nitrocellulose-coated slides spotted with increasing concentrations of purified BRD4 protein. We detected a robust BRD4 compound **1** dose-dependent interaction with a purified protein complex of biotinylated pVHL, Elongin B and Elongin C (bio-VBC) (**Fig. 1a**). Using this protocol, we probed high content functional protein microarrays (ProtoArray^®^) which contained >9000 purified human proteins with bio-VBC combined with either DMSO, **1** or a pooled mixture^13^ of three previously published VHL-binding molecules: compounds **2**^14^, **3**^15^ and **4** (**Table 1**). Notably, **1-4** all bind to pVHL with similar affinities (**Table 1**). To our surprise, we observed a strong signal for the spots corresponding to cysteine deoxygenase 1 (CDO1) for the compound pool (**2,3,4**) but not for **1** only (**Fig. 1b**). This result suggested that one or more of the compounds in the pooled mixture was responsible for inducing the bio-VBC-CDO1 interaction. To test this, full-length human CDO1 was expressed and purified from *E. coli* and tested for compound-induced interaction with bio-VBC using surface plasmon resonance (SPR) whereby bio-VBC was immobilized on a streptavidin chip and a titration of purified CDO1 was profiled over the surface in the presence of **2**, **3**, or **4 (Fig. 1c)**. No interaction of CDO1 with immobilized bio-VBC was observed with a titration of up to 4 µM CDO1. In contrast, addition of 10 µM **4** to CDO1 brought about saturable binding with bio-VBC with a calculated K_D_ of ∼0.82 µM. Neither 10 µM **2** nor 10 µM **3** led to a robust interaction of CDO1 with bio-VBC by SPR. To corroborate these results, we assessed **4**-dependent complex formation between VHL and CDO1 using TR-FRET and observed a **4** dose-dependent signal with a AC_50_/A_max_ of ∼111 nM/209 (**Table 1/Supplemental Fig. 1a, 1b**). We also examined **4** dependent ternary complex formation by 2D-NMR (**Fig. 1d**). NMR signals of VHL selectively methyl-group labeled on isoleucine, leucine and valine side chains were observed by ^13^C-^1^H correlation. In the presence of **4**, significant chemical shift perturbations were observed for a subset of methyl peaks relative to no compound. Subsequently, addition of CDO1 led to further chemical shift perturbations as well as a broadening of the VHL signals, further strengthening that CDO1 interacts with the VHL-compound **4** complex. In contrast, when CDO1 was added to VHL in the absence of **4**, no significant chemical shift perturbations of VHL signals were observed, demonstrating that interaction of VHL and CDO1 is **4**-dependent. Collectively, these results suggest that **4** acts as a molecular glue to recruit CDO1 to VHL through enabling *de novo* protein-protein interactions. While these results suggested that CDO1 is recruited to VHL by **4**, they did not exclude the possibility that **4** binds both CDO1 and VHL independently of one another. To address if **4** has any measurable affinity for CDO1 in the absence of VHL, we inverted the experimental SPR setup and immobilized biotinylated CDO1 (bio-CDO1) on a streptavidin chip. In this SPR configuration we did not observe interaction between bio-CDO1 and **4** with a titration of up to 10 µM in the absence of VHL (data not shown). We also did not observe interaction between bio-CDO1 and non-biotinylated VBC (**Fig. 1e**), consistent with our NMR observations. In contrast, when non-biotinylated VBC was added together with **4**, a robust interaction with immobilized bio-CDO1 was observed with a calculated K_D_ of ∼0.49 µM (**Fig. 1e**).

**Figure 1.**
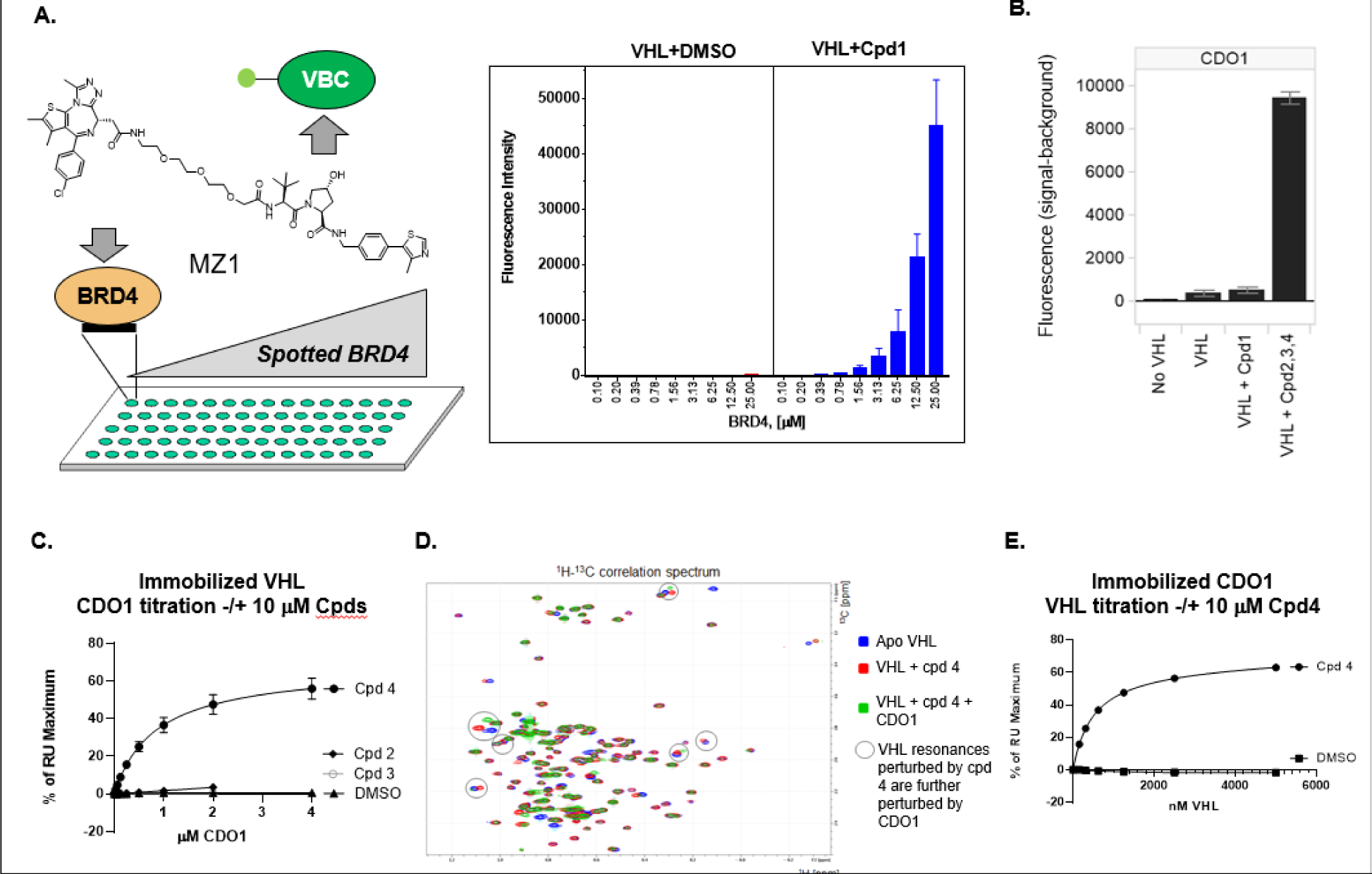
CDO1 is recruited to VHL by compound 4, a molecular glue. A. Cpd 1 recruits BRD4 to a bio-VBC probe on microarrays spotted with a titration of BRD4 protein. B. CDO1 is detected on Protoarrays^®^ when probed with bio-VBC and a pool of compounds 2-4. C. SPR with bio-VBC immobilized on a streptavidin chip and profiled with a titration of CDO1 up to 4 µM in the presence of 10 µM compound 2 or 3 or 4. D. 2D-NMR analysis of VHL, VHL + compound 4, or VHL + compound 4 + CDO1. E. SPR with bio-CDO1 immobilized on a streptavidin chip and profiled with a titration of VHL up to 5 µM in the presence of DMSO or 10 µM compound 4.

### Compound and VHL-dependent proteasomal degradation of CDO1

To test if **4** can recruit CDO1 to VHL in cells, we used a NanoBiT (Promega) split-luciferase complementation assay (**Fig. 2a**). Constructs expressing VHL-LgBiT and SmBiT-CDO1 were transiently co-transfected into HEK293T cells. In the presence of **4**, we observed robust dose-dependent signal consistent with ternary complex formation with a AC_50_/A_max_ of ∼334 nM/578. Consistent with the biophysical assays, **3** did not lead to formation of a VHL-cpd-CDO1 ternary complex by NanoBiT (**Table 1**). Intriguingly, **2** did show some recruitment with an AC_50_/A_max_ of ∼731 nM/232. We then asked if compound-dependent recruitment of CDO1 to VHL leads to degradation of CDO1 in cells using a HiBiT split-luciferase degradation assay. HiBiT-tagged CDO1 was transiently transfected into HEK293T cells, which express VHL, and cells were treated with **4** in dose response for 19 hours. We observed data consistent with **4**-induced CDO1 degradation (absDC_50_ of 349 nM/77% degradation) (**Fig. 2b/Table 1**). No degradation was observed with **2** or **3** (**Table 1**). Although these results are consistent with VHL-**4**-dependent CDO1 degradation, we wanted to further address VHL-dependence in a VHL-negative renal cell adenocarcinoma cell line (786-0)^16,17^. To test CDO1 degradation in 786-0 cells, CDO1 was virally transduced because we could not detect endogenous CDO1 in 786-0 by western blot (data not shown). The CDO1-expressing 786-0 cells were subsequently transduced with either wild-type VHL or an empty expression vector yielding 786-0^CDO1+/VHL+^ and 786-0^CDO1+/VHL-^ cell lines respectively. As expected, endogenous HIF complex subunit Hif-2, which is regulated by VHL^1^, was degraded in 786-0^CDO1+/VHL+^ cells but not in 786-0^CDO1+/VHL-^ cells (**Fig. 2c**). After 786-0^CDO1+/VHL+^ and 786-0^CDO1+/VHL-^ cells were treated with DMSO or 10 µM of **4** for 16 hours, we observed degradation of CDO1 in 786-0^CDO1+/VHL+^ cells but not in 786-0^CDO1+/VHL-^ (**Fig. 2c**). These data validate that degradation of CDO1 by **4** is dependent on VHL. As VHL is a component of a multi-subunit cullin-ring E3 ligase complex which requires post-translational neddylation of CUL by the NEDD8-activating enzyme NAE1 to effect target ubiquitination and degradation, we added the NAE1 inhibitor^18^ (MLN4924) and showed that 786-0^CDO1+/VHL+^ cells treated with **4** in the presence of 1 µM MLN4924 or 1 µM of the proteasome inhibitor MG132 blocked compound-induced degradation (**Fig. 2d**). Collectively, these data confirm that **4** is a VHL dependent molecular glue degrader for CDO1.

**Figure 2.**
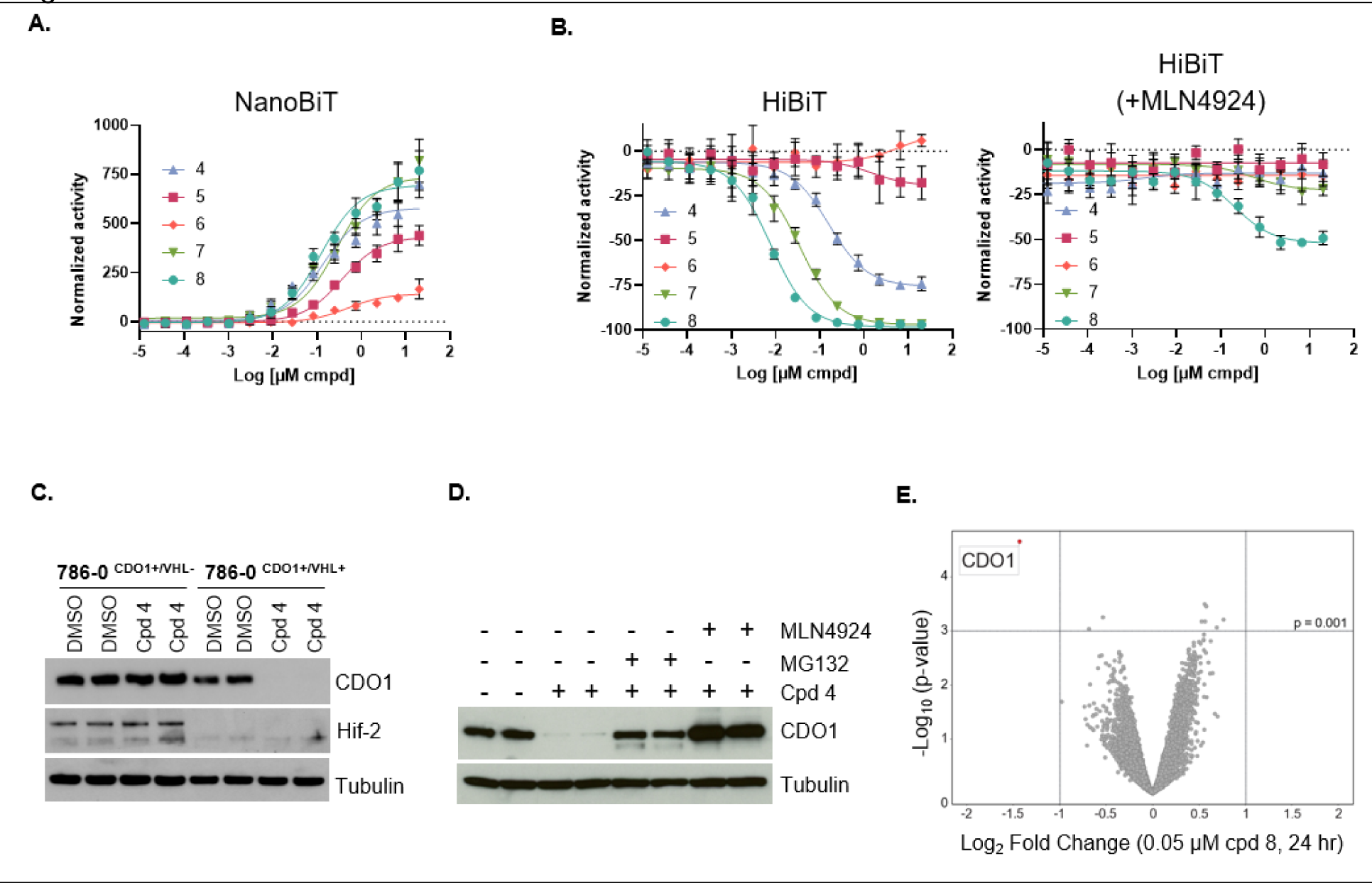
Recruitment of CDO1 to VHL by ligands based on cpd 4 leads to CDO1 degradation. A. Cellular recruitment of CDO1 to VHL by NanoBiT. VHL-LgBiT and CDO1-SmBiT plasmids were co-transfected into HEK293T cells in the presence of DMSO or a dose-response of VHL ligands 4-8. Normalized activity is percent of DMSO control. B. HiBiT degradation assays. HEK293T cells were transfected with CDO1-HiBiT and then subject to a dose-response with VHL ligands, without or with 1 µM MLN4924 treatment for 8 hrs addition of VHL ligands 4-8. After 16 hours, cells were lysed in HiBiT lysis buffer and read on a luminescence reader. Normalized activity is percent signal change relative to DMSO control. C. Western blot of CDO1 degradation in 786-0^CDO1+/VHL-^ and 786-0^CDO1+/VHL+^ cells. Cells were treated with DMSO or 10 µM compound 4 for 16 hours before harvesting for Western blotting. D. Western blot of CDO1 degradation in 786-0^CDO1+/VHL+^ cells. Cells were treated with 10 µM compound 4 for 16 hours with or without prior treatment with 1 µM MG132 or 1 µM MLN2934 before harvesting. E. Quantitative proteomics profiling of Huh-7 cells treated for 24Lhours with 0.05 µM of compound 8 or DMSO. Volcano plot shows downregulation of endogenous CDO1 upon compound 8 treatment compared to DMSO (protein FDR < 1%, nL=L3 biological replicates per treatment). Log_2_(fold change) difference between means of treated vs. DMSO plotted against p-values calculated using Limma. Lines in the plot indicate significant cutoffs: p-value < 0.001 and absolute [Log_2_(fold change)] > 1.0.

### Optimization of CDO1 recruitment and degradation

To understand the structure-activity relationship of **4** as a glue for VHL and CDO1, we synthesized and profiled a small number of analogs. Replacement of the t-butyl group on the left-hand side of **4** with an isopropyl group yielded compound **5** and led to a drastic decrease of CDO1 degradation to only 18% which was associated with decreased recruitment of CDO1 by TR-FRET, SPR and NanoBiT (**Table 1**, **Fig. 2a/b**). Given that **5** has similar binding affinity to VHL, this suggested that the additional methyl group could be important for CDO1 recruitment. Exploration of the right-hand side showed that while the addition of a methyl group adjacent to the benzylic position, yielding compound **6,** slightly improved VHL binding, this modification drastically reduced CDO1 recruitment and degradation (**Table 1**, **Fig. 2a/b**). Conversion of the phenyl ring to the larger naphthyl ring yielding compound **7** did improve both biochemical/cellular recruitment and degradation relative to **4** (**Table 1**, **Fig. 2a/b**). Finally, replacement of the methyl-thiazole ring of **7** to a pyrazole, led to the creation of compound **8**, which recruited CDO1 with the highest affinity of 0.10 μM and was also the most potent degrader of CDO1 (absDC_50_ of 8nM/98% degradation) (**Table 1**, **Fig. 2a/b**). For all compounds that degraded CDO1, MLN4924 blocked or strongly reduced degradation (**Fig. 2b**). To eliminate the possibility that these VHL ligands were causing cell death, we performed a Cell Titer Glo cell viability assay with **5** and **8**, representing very weak and potent degraders respectively (**Supplemental Fig. 1c**). Treatment with either ligand did not have any detectable effect on cell viability. Interestingly, unlike the biochemical recruitment experiments, the cellular NanoBiT assay was able to detect some recruitment for the weaker **5**, but it did not significantly distinguish between **4**, **7** and **8**. In addition, **2** had significant (albeit weaker than **4**) activity in the cellular recruitment assay but did not have activity in biochemical recruitment or CDO1 degradation assays. Thus, the biochemical experiments were better at differentiating compounds capable of inducing CDO1 degradation.

To gain a better understanding of the selectivity of degrader **8**, we treated Huh-7 cells with 50 nM **8** for 24 hrs and assessed the proteomic profiles using mass-spectrometry. Robust and selective degradation of endogenous CDO1 was observed with treatment with **8**, among the 9,478 quantified proteins, with a >3.5-fold change normalized to the DMSO control, p-value <1.0E-003 and an overall degradation of 62%. CDO1 was identified with four unique quantified peptides in biological triplicate confirming its significant modulation. (**Fig. 2e**).

### Prediction of the VHL recruitment interface on CDO1

To address the molecular basis of **4**-mediated recruitment of CDO1 to VHL, we applied *in silico* protein-protein docking to model potential VHL-**4**-CDO1 ternary complexes. From this first docking step, seventeen complexes having the t-butyl group of **4** within the protein-protein interface (PPI) were carried forward. The complexes were clustered by ligand location on CDO1 resulting in six plausible ligand binding CDO1 regions (**Supplemental Fig. 2a**). Local protein-protein docking was performed around these six regions and the poses were scored and clustered resulting in eight possible complexes. These were further assessed by MD simulations^19^ followed by water solvation analysis with WATMD^20,21^ (**Supplemental Fig. 2b**) and the water structure at the PPI was determined. The conformations were ranked by the degree of de-wetting at the PPI as seen in these calculated water structures, with one highly plausible contact area (**Supplemental Fig. 2c**). In that region, the ligand makes contact with a helical region around W47 as well as in the spatially (but not in sequence) proximal P150 region.

In a parallel effort, we aimed to use CDO1 species variation with the goal to define key regions important for **4** recruitment. To do this, we expressed and purified CDO1 proteins from 11 species followed by testing by SPR for ternary complex formation (**Supplemental Fig. 3a)**. Interestingly, only the CDO1 protein of *Cyprinus carpio* (common carp) failed to be recruited to VHL (**Fig. 3a**). A sequence alignment of human and carp CDO1 revealed regions of sequence dissimilarity which we arbitrarily clustered into four regions, annotated by their coordinates on human CDO1: R1(Q3-E27), R2(A35-Q65), R3(N90-R126), and R4 (I145-L197) (**Fig. 3b**). We then designed, expressed, and purified eight chimeric human-carp CDO1 proteins in which each of the four sequence dissimilarity clusters were swapped, and tested the chimeric human-carp CDO1 proteins for ternary complex formation by SPR (**Supplemental Fig. 3b**). Carp region R1 swapped into human CDO1 (hum-carp 1) did not modulate recruitment and exchanging human R1 into carp did not restore recruitment (hum-carp 5). This data suggested that R1(Q3-E27) was most likely not important for recruitment. Additional chimeras in which regions R2, R3 or R4 were swapped were all associated with decreased recruitment of CDO1 suggesting each of these regions, R2(A35-Q65), R3(N90-R126) and R4 (I145-L197) are likely important for recruitment (**Supplemental Fig. 3b**). We combined these chimeric protein results with the contact information from protein-protein docking poses around the most plausible PPI contact area to arrive at six more narrowly defined regions within the original proposed regions R2, R3 and R4. The refined regions are annotated as R2’(D43-A48), R3’(M97-N101), R3’’(S114-E122), R3’’’(R126), R4’(P150) and R4’’(D162-N200) (**Fig. 3b/Supplemental Fig. 3c**). To test which of these refined regions were important for recruitment of CDO1, 12 additional human-carp chimeric proteins were produced and tested by SPR for **4**-mediated recruitment to VHL (**Fig. 3c**). Hum-carp 15, which contains human regions R2’(D43-A48), R3’(M97-N101), R3’’(S114-E122), R3’’’(R126), R4’(P150) and R4’’(D162-N200) introduced into Carp CDO1, was robustly recruited. In contrast, hum-carp 9, which contains carp regions R2’(K43-M48), R3’(L97-Q101), R3’’(Q114-Q123), R3’’’(L127), R4’(T151) and R4’’(Q163-201) introduced into human CDO1, was not recruited. Taken together, these data argue that some or all of the human regions R2’(D43-A48), R3’(M97-N101), R3’’(S114-E122), R3’’’(R126), R4’(P150) and R4’’(D162-N200) are important for recruitment. Hum-carp 18, which contains only human R2’(D43-A48) and human R4’(P150) within carp CDO1, was recruited just as well as human-carp 15 arguing that human R3’(M97-N101), human R3’’(S114-E122), human R3’’’(R126) and human R4’’(D162-N200) are not essential for recruitment. Hum-carp 12, which contains carp regions R2’(K43-M48) and R4’(T151) in human CDO1 was not recruited. These data support a critical role for human regions R2’(D43-A48) and R4’(P150) in recruitment of CDO1 to VHL bound to **4**. Thus, the initial recruitment interface proposed by *in silico* protein-protein docking is in agreement with the experimentally determined regions required for recruitment.

**Figure 3.**
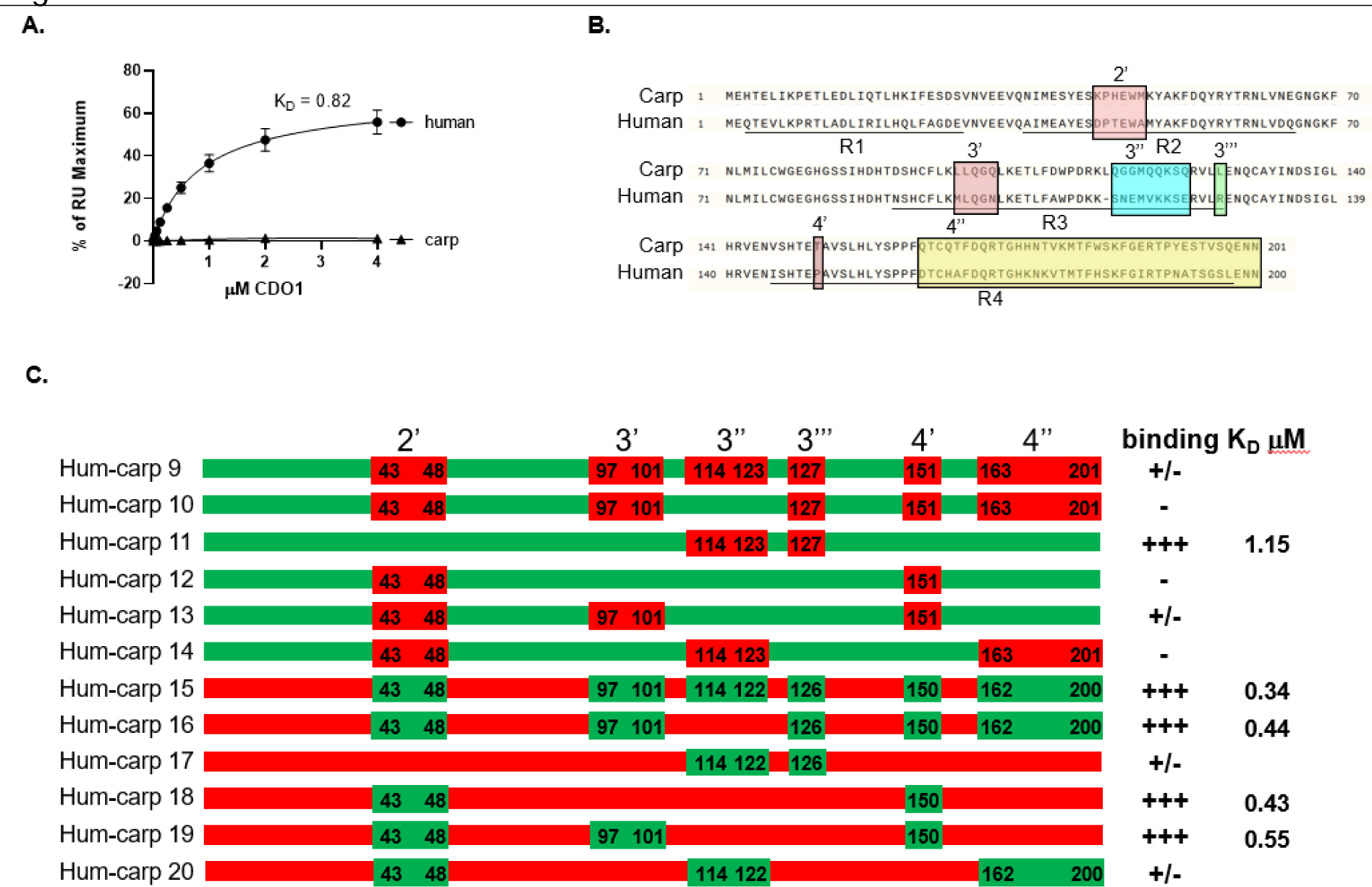
Recruitment of CDO1 species variants/chimeric proteins to VHL and proposed recruitment interface of CDO1. A. SPR for titration of either human or carp CDO1 to bio-VBC with 10 μM compound 4. B. Sequence alignment of human and carp CDO1 proteins. Regions of sequence differences clustered into several regions denoted as R1-R4 (underlined). Refined subregions within the R2-R4 clusters based on predicted protein-protein contacts are boxed and denoted as 2’, 3’, 3’’, 3’’, 4’ and 4’’. C. Schematic of refined human-carp chimeric proteins with human sequences in green and carp sequences in red. Relative incorporation into a ternary complex with bio-VBC in the presence of 10 μM compound 4 as measured by SPR is indicated by +/- signs and binding constants (K_D_) where calculable.

### X-ray structures for VHL-cpd-CDO1 ternary complexes

We then attempted to obtain X-ray crystal structures of complexes with VHL, CDO1 and with **4** or **8**. High protein complex concentration and low ionic strength in the final protein formulation were used to facilitate rapid crystallization by avoiding the slow oxidation of human CDO1 ferrous ion^22^. Crystals were obtained for both VBC-**4**-CDO1 and VBC-**8**-CDO1 complexes and structures were determined to 2.9 Å and 2.5 Å resolution respectively (**Supplemental Table 1**).

The structures reveal 1:1:1 complexes where **4** and **8** are sandwiched in between VHL and CDO1 (**Fig. 4a**). As expected, both compounds make similar side-chain/back bone polar interactions with VHL residues Y98/R107/H110/S111/H115. Of particular interest are the *de novo* VHL-CDO1 PPI interactions surrounding the glue-binding site, effectively occluding **4** and **8** from solvent **(Fig. 4b).** The interface is stabilized by multiple protein-protein interactions observed in both ternary complexes involving 12 and 14 VHL and CDO1 residues respectively **(Supplemental Fig. 4a-c)**. As this ring of de novo PPI contacts serves to occlude water molecules from the glue-binding site, it is reminiscent of the “O-ring” typically found at protein-protein interfaces^23^ **(Supplemental Fig. 4c)**.

**Figure 4.**
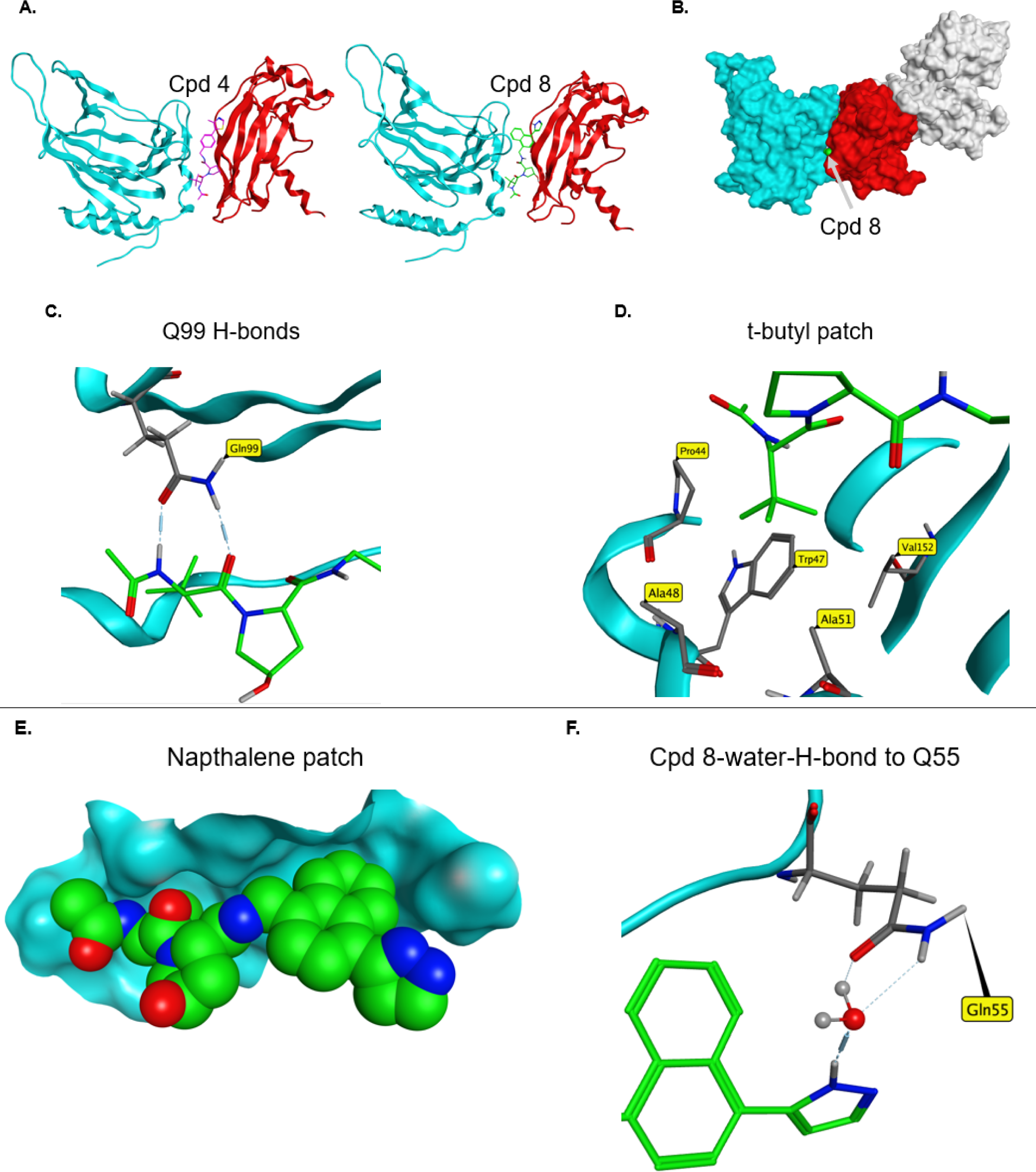
Molecular glue interactions for VHL-cpd-CDO1 ternary complexes. A. X-ray ternary structures for compound 4 (purple, PDB ID 8VLB) or compound 8 (green, PDB ID 8VL9) showing 1:1:1 stoichiometry with CDO1 (blue) and VHL (red). B. Surface display showing complete engulfment of compound 8 (green) between CDO1 (blue) and VHL (red). Elongin B/C is shown in grey. C. Hydrogen bonds formed between compound 8 with Q99 of CDO1. D. Key CDO1 residues involved in hydrophobic interactions with the t-butyl group of compound 8. E. The naphthalene of compound 8 (green/space-filling view) fills a cavity bounded by several residues F53/D54/G78/P150, with surface map of CDO1 shown in blue. F. The pyrazole group of compound 8 makes a water-mediated H-bond with Q55.

The small molecule interactions with CDO1 are in alignment with the regions predicted by the modeling efforts and experimentally determined by assessing CDO1 proteins from various species. Glutamine 99 of CDO1 makes 2 hydrogen bonds with the left-hand side of both **4** and **8 (Fig. 4c)**. Noteworthy is that Q99 is conserved in both the human and carp CDO1 sequences. To test the hypothesis that CDO1 Q99 is making critical H-bond contacts, Q99 was mutated to asparagine (Q99N) and tested for recruitment by SPR (**Supplemental Fig. 5a**). Relative to wild type CDO1, **4**-mediated recruitment of CDO1(Q99N) to VHL was reduced by greater than 40-fold with a K_D_ >40 μM. Q99N also reduced recruitment of CDO1 with **8** by 47-fold and a K_D_ of ∼6.1 μM. CDO1(Q99N) was also tested for recruitment to VHL by TR-FRET (**Supplemental Fig. 5b**). Relative to wild type CDO1, CDO1(Q99N) reduced the recruitment AC_50_ in presence of **4** by ∼158-fold with an AC_50_ of ∼17.4 μM, while reducing the recruitment AC_50_ in presence of **8** by ∼29-fold with an AC_50_ of ∼ 0.87 μM, respectively. CDO1(Q99N) was also tested for its ability to get degraded in cells (**Supplemental Fig. 5c**). Transiently expressed HiBiT-tagged CDO1(Q99N) showed no apparent degradation in the presence of either **4** or **8**. Taken together, these results are consistent with CDO1(Q99) making critical contacts with both **4** and **8** which is essential for robust recruitment and degradation in cells. The X-ray structure also reveals that the t-butyl group on the left-hand side of the molecule fits nicely into a CDO1 hydrophobic patch formed by residues from P44, W47, A48, A51, and V152 (**Fig. 4d**) consistent with its importance determined in earlier experiments. The naphthalene group on the right-hand side fits into a pocket bounded by residues F53, D54, Q55, G78, and P150 (**Fig. 4e**). The improved affinity of **8** relative to **4** is likely due to the naphthalene group better filling of the hydrophobic cavity relative to the phenyl group in **4** and also from a water-mediated H-bonding interaction of compound 8 with Q55 (**Fig. 4f**). The VHL-small molecule glue effectively recognizes a composite interface formed by discontinuous regions of CDO1.

## Discussion

The reports that lenalidomide recruits CRBN to degrade IKZF1/3^24,25^ have spurred a rapid increase in the study of molecular glue degraders for the development of novel therapeutics. These initial mechanism of action studies have led to the expansion of the CRBN neosubstrate scope to include therapeutically relevant yet previously undruggable targets such as GSPT1^26,27^ and IKZF2^27^ enabled in part by the identification of a simple beta-hairpin degron^28^ as well as collections of CRBN ligands. Analogously, studies of the sulfonamide indisulam have also revealed that it functions as a DCAF15-based glue of RBM39^29–33^. The studies of target focused libraries have led to the identification of DCAF16-based glue degraders of BET bromodomains^34^ and CDK12 ligands that recruit CDK12 to DDB1, leading to the degradation of CDK12’s substrate, cyclin K^35^, but the potential target scope of this approach is somewhat limited by the need for chemical matter binding to relevant targets.

Rather than rely on phenotypic screening, we sought to use protein arrays to identify neosubstrates for VHL using previously developed small molecules that have been used as handles for bifunctional degraders, but without any known molecular glue activity. Some advantages of using protein arrays for neosubstrate identification are the relative normalized protein concentrations across the array compared to the wide range observed in cells and the ability to easily control assay parameters. Consequently, the method allows for the detection of weak molecular glue interactions that could be easily missed in cellular screens for protein recruitment and/or degradation. We were fortunate to identify the VHL molecular glue target, CDO1, on the protein array because CDO1 is an example of a protein whose expression is restricted to certain tissues, primarily lung, duodenum, small intestine, liver and appendix, and could easily be missed in proteomic screens. To date, only a few companies have produced high content functional protein arrays that can be used for E3 ligase neosubstrate identification as the process for making thousands of high-quality full-length proteins that are functional on surfaces is not trivial.

CDO1 is an enzyme that catabolizes cysteine by adding a molecular oxygen to the sulfur of cysteine to form cysteine sulfinic acid^36^. In the liver, CDO1 functions to remove excess cysteine obtained from the diet as high levels of cysteine are cytotoxic to neurons and cells^37^. CDO1 also appears to act as a tumor suppressor in numerous cancers, as CDO1 expression is repressed by promoter methylation and this repression is linked to poor prognosis^38^. CDO1 protein stability is regulated by cysteine levels, being unstable in low cysteine and stable at high cysteine levels^39^ and this regulation by cysteine was shown to be proteasome dependent but the E3 ligase responsible has not yet been identified. We confirmed that CDO1 regulation by cysteine is still intact in cells that lack VHL, suggesting that VHL is not the cysteine regulating ligase and that CDO1 is a true neosubstrate for the discovered VHL glues (**Supplemental Fig. 6**). Lastly, we could not identify a disease context in which VHL ligase compound-dependent depletion of CDO1 would be beneficial.

An examination of the degron in CDO1 reveals that it is bipartite, meaning key contact regions with VHL are localized to different regions of CDO1. Because **4** and **8** have no measurable affinity to CDO1, each bring about the formation of a high affinity/cooperative ternary complex. When we examined the proteome by searching structural databases for proteins that would have a similar degron, we did not find anything with high similarity. The unique nature of the CDO1 degron is further supported by our inability to observe degradation of any other cellular protein in proteomic analyses across several cell lines with either **4** or **8**. However, we do know that VHL molecular glues are extendable to other targets as we have been able to show from additional profiling experiments with VHL chemical matter distinct from the small molecules in this manuscript (data not shown), that VHL molecular glue degraders for additional novel targets with high cooperativity are achievable, and will enable investigations into perturbing targets/classes of proteins that are currently not druggable using CRBN IMIDs or bi-functionals.

Given the clinical relevance of glue degraders such as thalidomide, lenalidomide and pomalidomide, as well as the recent expansion of therapeutically relevant neosubstrates for CRBN, there is an acute need for additional techniques to identify neosubstrates. Mason et al. have addressed this challenge by generating a ∼1 million compound, diverse hydroxy-proline scaffold directed DNA-encoded library (DEL) chemical library which was employed in target screens to identify candidate molecular glues for the BRD4 bromdomain^40^. Off-DNA synthesis and testing of the hits which were enriched for BRD4 only in the presence of VHL revealed these molecules indeed recruit VHL to BRD4 bromodomain and also bring about VHL dependent degradation of endogenous BRD4 in cells. In this approach only one type of library was used, but an aspiration of this technology is that by employing highly diverse libraries with multiple presenter chemical scaffolds, additional novel molecular glues will be identified. Molecular glues are without a doubt a very exciting area of research and novel approaches to designing and identifying chemical matter with the capability to target undrugged proteins will continue to be needed to advance both chemical probes and compounds in the clinic.

## Author contributions

Antonin Tutter performed the protein array experiments that led to the identification of CDO1, expressed and purified the wild-type, Q99N, species-specific and chimeric CDO1 proteins, and performed the TR-FRET, NanoBiT, HiBiT and CTG assays. Dennis Buckley, Veronique Darsigny and Aleem Fazal designed and synthesized VHL ligands. Xiaolei Ma and Wei Shu solved the X-ray ternary complex structures. Andrei Golosov performed the protein-protein docking analysis and Daniel McKay performed the WATMD analysis for ternary complex interface predictions which were used in chimera designs for interface mapping. Rohan Beckwith proposed the idea of using protein arrays to find molecular glue targets for VHL. Jonathan Solomon conducted the compound **4** experiment in 786-0^CDO1+/VHL-^ and 786-0^CDO1+/VHL+^ cell lines. Pasupuleti Rao ran the MLN4924 and MG132 experiment in 786-0^CDO1+/VHL-^ and 786-0^CDO1+/VHL+^ cell lines. Lei Xu generated the 786-0^CDO1+/VHL-^ and 786-0^CDO1+/VHL+^ cell lines. Andreas Lingel performed the NMR experiments. Charles Wartchow performed the SPR assays to measure compound affinities with VHL. Jennifer Cobb and Amanda Hachey conducted the proteomics experiments. Dustin Dovala expressed and purified biotinylated and unbiotinylated pVHL complexes. Gregory A. Michaud performed the SPR assays for ternary complex formation and is the corresponding author.

## Methods

### Protein expression/purification

Biotinylated pVHL-Elongin B-Elongin C (bio-VBC): VHL(54-213) with an N-terminal 8xHistidine-3C-AVI tag was synthesized in a single vector along with Elongin B and Elongin C(17-112). A single T7 promoter drove the expression of the tricistron, and each gene individually had a 5′ ribosome binding site and epsilon translational enhancer element. The plasmid was used for transformation of C41(DE3) *E. coli* cells using standard techniques. Cells were grown under standard growth conditions in terrific broth to mid-log phase, at which point they were induced with IPTG and left to grow overnight at 20°C. The cells were then harvested via centrifugation and resuspended in lysis buffer (25 mM Tris pH 7.5, 200 mM NaCl, 10 mM imidazole, and 1 mM TCEP), whereupon they were lysed via cell homogenizer (3 passes at 18,000 psi). The lysate was clarified by ultracentrifugation (160,000 x g for 2 hours). Cleared lysate was subjected to immobilized metal affinity chromatography (IMAC) using nickel-conjugated NTA resin. The resin was initially washed with 5 column volumes (CV) of lysis buffer, followed by 5 CV of wash buffer (25 mM Tris pH 7.5, 200 mM NaCl, 40 mM imidazole, and 1 mM TCEP), followed by elution with 5 CV of elution buffer (25 mM Tris pH 7.5, 200 mM NaCl, 500 mM imidazole, and 1 mM TCEP). The IMAC eluate was subject to biotinylation overnight at 4°C via the addition of 200 ug/mL biotin, 10 mM ATP, 10 mM MgSO4, 1 mM TCEP and 500 μg BirA ligase. The progress of the reaction was monitored by ESI-LC/MS and went to completion. Biotinylated VHL complex was then subjected to cleavage by HRV 3C protease while dialyzing against dialysis buffer (50 mM Tris pH 8.0, 100 mM NaCl, 1 mM TCEP). Cleavage was monitored by LC/MS and went to completion over 2 hours. The dialyzed protein was diluted 2:3 with water and loaded onto a Q HP column at 4 mL/min on an AKTA FPLC. The protein was eluted with a linear gradient from 100% IEX Buffer A (50 mM Tris pH 8.0, 50 mM NaCl) to 100% IEX Buffer B (50 mM Tris pH 8.0, 500 mM NaCl) over 8 CV at 4 mL/min. All fractions from the peak were combined. The protein was then subjected to reverse IMAC purification by flowing through 5 mL of Ni-NTA resin. The flow-through was collected and concentrated with a centricon 3,000 MWCO concentrator. Lastly, the protein was subjected to size exclusion chromatography utilizing a Superdex 200 16/60 column, pre-equilibrated with SEC Buffer (25 mM Tris pH 7.5, 150 mM NaCl, 1 mM TCEP). Protein was loaded and run at 1 mL/min. The complex eluted as a single peak, with lower MW shoulder. All fractions from this peak were combined, concentrated, and aliquoted. Total yield was approximately 6 mg/L of culture. For crystallography, a similar construct omitting the AVI tag and using a truncation of Elongin B (1-104) was utilized. Purification proceeded similarly, with the exclusion of the biotinylation step.

CDO1: For crystallography, a plasmid encoding wildtype CDO1 containing an N-terminal TEV-cleavable poly-histidine tag was used to transform BL21(DE3) cells. Cells were grown under standard conditions in terrific broth to mid-log phase, at which point the temperature was decreased to 18°C and IPTG was added to the cultures to a concentration of 1 mM. Growth continued for 19 hours. Cells were harvested and the protein was subjected to lysis and IMAC as done for VHL. The protein was then supplemented with 0.5 mM EDTA and cleaved with TEV protease overnight while dialyzing in dialysis buffer. This was followed by reverse IMAC and size exclusion chromatography as previously described. Total yield was approximately 30 mg/L of culture. For TR-FRET and SPR, all versions of CDO1 were expressed as an N-terminal 6His fusion as described above, with the exception that the 6-His tag was not cleaved after the initial IMAC and SEC purification. For bio-CDO1, a plasmid encoding wildtype CDO1 containing an N-terminal 6Histidine-TEV-AVI tag was used to express and purify protein as above for TR-FRET and SPR, with the exception that the protein was biotinylated and TEV-cleaved similarly as for VHL after IMAC and prior to SEC purification.

### Protein arrays

To generate the protein arrays spotted with a BRD4 titration, the indicated concentrations of purified 6His-tagged BRD4 were spotted onto blank nitrocellulose-coated glass slides (Grace Bio-Labs PATH protein microarray slides, Cat# 805020) using a SpotBot 3 microarrayer (ArrayIt cat# SPA3PRO). The arrays were stored in sealed array holders at −20°C for at least 72 hours before use. For probing the spotted BRD4 arrays and the Invitrogen ProtoArrays (Invitrogen cat# PAH0525101) with bio-VBC and compounds, a protocol closely based on the Invitrogen ProtoArray protocol (https://tools.thermofisher.com/content/sfs/manuals/protoarray_applicationsguide_man.pdf) was used. All steps were performed at 4°C unless otherwise noted. Arrays were first incubated in 5 mL of block buffer (50 mM HEPES pH 7.5, 200 mM NaCl, 0.08% Triton X-100, 25% glycerol, 20 mM reduced glutathione, 1 mM DTT, and 1X synthetic block [Invitrogen cat# PA017]) for one hour with gentle rocking, in a four-chamber microarray incubation tray (Sarstedt, Cat. no. 94.6077.307). Typically, three or four arrays were used in a single experiment. During the blocking step, bio-VBC probe was prepared by diluting bio-VBC to 10 µM in 1X block buffer along with compounds or DMSO. For the probe containing compounds 2-4, each compound was added at 3.33 µM for a 10 µM total compound pool concentration in 1.6% final DMSO. For the probe containing compound 1, 1 was added at 3.33 µM in 1.6% final DMSO. After blocking, block buffer was aspirated from each chamber containing an array, and 120 µL of each bio-VBC probe was dispensed drop-wise directly onto the surface of the arrays and covered with a LifterSlip cover slip (Thermo cat# 25X60I-2-4789). The arrays were incubated with the probes for 90 minutes. The coverslips were removed using forceps, and the arrays were washed in 5 mL of wash buffer (1X PBS, 0.1% Tween-20, 1X synthetic block) for five minutes with gentle rocking. Streptavidin-Alexafluor 647 (Thermo cat# S-32357) was added to wash buffer at 1 µg/mL, and 5 mL was dispensed onto the arrays and incubated for 45 minutes with gentle rocking. Five additional washes were performed, 5 mL each for five minutes. The arrays were transferred to room temperature and dipped into room-temperature water in a 50 mL conical tube for one second to remove any residual components of wash buffer, then centrifuged at 200 x g for two minutes in the slide carrier provided with the Grace PATH arrays to remove any residual liquid. The dry slides were scanned on a GenePix 4000B slide scanner and the images were analyzed using ProtoArray Prospector software. The resulting data table was imported into Spotfire analysis software (Tibco) and Z-scores were calculated for each array. Hits were scored by filtering arrays probed with bio-VBC and compounds 2-4 for Z-scores >4, and filtering arrays probed with bio-VBC without compounds, bio-VBC with compound 1, or Alexafluor 647 alone, for Z-scores <1. Using this filtering scheme across all four arrays resulted in eight hits including CDO1.

### Surface plasmon resonance

Biotinylated VBC complex (bio-VBC) or biotinylated CDO1 (bio-CDO1) was immobilized on a streptavidin chip using a Biacore T200 or a Biacore 8K (GE Healthcare). To determine the kinetics-affinity of the small molecules for proteins, small molecules were diluted in DMSO such that final concentration gradients in DMSO were 50X. 3.1 µL of the small molecules were then added to 150 µL of buffer (50 mM Tris pH 8.0/150 mM NaCl/0.01% Tween 20/1mM EDTA/1 mM DTT) in a 96/384 well plates and mixed using a Biomek FX. The small molecule solution gradients were then injected at 45 µL/min for 120 seconds contact time and dissociation time was 180 seconds in running buffer (50 mM Tris pH 8.0/150 mM NaCl/0.01% Tween 20/1mM EDTA/1 mM DTT/2% DMSO). The kinetics data was fit to a 1:1 binding model to measure the association rate ka (1/Ms), the dissociation rate kd (1/s) and the affinity K_D_(M). To determine the apparent kinetics-affinity for ternary complex formation for small molecules bound to VHL with CDO1, small molecules were diluted in DMSO such that final concentration in DMSO was 500 μM. his^6^-CDO1(full-length) was diluted in buffer (50 mM Tris pH 8.0/150 mM NaCl/0.01% Tween 20/1mM EDTA/1 mM DTT) to prepare concentration gradient (8 concentrations/2 fold dilutions). 3.1 µL of the 500 μM small molecules was added to 150 µL of the CDO1 samples in a 96/384 well plates. The CDO1 solution gradients were then injected at 45 uL/min for 120 seconds contact time and dissociation time was 180 seconds in running buffer (50 mM Tris pH 8.0/150 mM NaCl/0.01% Tween 20/1mM EDTA/1 mM DTT/2% DMSO). The data was fit to a steady-state model to measure the apparent affinity K_D_ (M) for ternary complex formation.

### TR-FRET

Ternary complex TR-FRET assays were performed with a mixture of 100 nM purified recombinant VHL protein complex consisting of N-terminally Avi-tagged and biotinylated VHL, untagged Elongin-B and untagged Elongin-C (bio-VBC), with 100nM C-terminally 6His-tagged CDO1, 1nM anti-6His-Europium (ThermoFisher PV5596) and 50nM SA-APC (ThermoFisher SA1005) in assay buffer (100 mM NaCl, 50 mM Tris pH 7.5, 0.1% Pluronic F127). The mixture was dispensed into 1536-well plates (Greiner # 782075), 5 µL/well using a Multidrop Combi (ThermoFisher). 50 nL of compounds in DMSO was added to target wells using an acoustic liquid handler (Labcyte Echo 500). The plates were incubated at room temperature for 60 minutes, and then read on a plate reader (Perkin Elmer Envision) using a TR-FRET protocol (Europium donor exc: 340/80 nm, ems: 615 nm; APC acceptor ems: 665 nm). The TR-FRET ratio was calculated as the ratio: signal 520 nm/signal 492 nm. The data was normalized using the equation 100*(well value-NC)/NC, where NC is the neutral control well (DMSO). AC50 values were calculated from curve fits generated by Spotfire analysis software (Tibco).

### NMR

NMR experiments were performed at 300 K on a Bruker 600 MHz Avance III spectrometer equipped with a 5 mm QCI-F z-gradient cryoprobe. Samples were prepared in phosphate buffered saline (PBS) and 5% D2O (v/v) and were assessed in 3 mm NMR tubes filled with 160 µL sample volume. All samples contained 50 µM VCB complex which was selectively ^1^H/^13^C methyl labeled on isoleucine, leucine and valine side chains in a uniformly 15N labeled and perdeuterated background^41^. Compound 4 was solubilized in DMSO-*d_6_* and added to a final concentration of 75 µM, whereas CDO1 was added to a final concentration of 50 µM. 2D NMR spectra were recorded with the [^13^C, ^1^H]-HMQC SOFAST experiment^42^ with 12 transients, 160 indirect points (in ^13^C dimension), and an ^1^H acquisition time of 142 ms.

### Cell lines and cell-based assays: 786-0^CDO1+/VHL-^ and 786-0^CDO1+/VHL+^ cell lines

pENTR-m.cherry and pENTR221-VHL (NM_198156) were Gateway LR cloned into the destination vector pLENTI4/V5 DEST. Each resultant plasmid was co-transfected with ViraPower™ Lentiviral Packaging Mix (ThermoFisher # K496000) into 293FT cells to make virus particles. Virus particles were filtered through a 0.4 micron filter and used to infect 786-0 cells as described previously^17^. Selection was performed with 500 µg/ml of <u>Zeocin</u> for two weeks to obtain 786-0^m.cherry/VHL-^ and 786-0^VHL+^ pooled cell lines. Both cell lines were then transduced with lentivirus harboring CDO1 (NM_001801) using the same technique as for VHL and m.cherry, but with 400 µg/mL G418 and 500 µg/ml of Zeocin dual selection for two weeks to obtain 786-0^CDO1+/VHL-^ and 786-0^CDO1+VHL+^ cell lines.

### Westerns

For treatment of 786-0^CDO1+/VHL-^ and 786-0^CDO1+VHL+^ cell lines with compounds, cells were resuspended in DMEM (Gibco # 21013-024) supplemented with 10% FBS (Gibco # 26400-044) and 1.25 mM cysteine and plated at 0.4 x 10 ^6^/mL in six-well plates. The next day, cells were treated with 30 µM compound 4 or DMSO (Figure 2C), or 10 µM compound 4 and either DMSO, 1 µM MG132 or 1 µM MLN4924 (Figure 2D). Cells were harvested 16 hours later, washed once with PBS, transferred to ice and lysed with 120 µL per well of IP buffer (ThermoFisher # 87787) containing protease inhibitor cocktail (Sigma # 11836170001). Cells were scraped and transferred to microfuge tubes and centrifuged at 13k x g for 5 minutes at 4°C. 10 uL of lysates was loaded on 4-12% NuPAGE Bis-Tris gels (Thermo NP0303) in MES buffer (Thermo NP0060). Proteins were transferred from gels to nitrocellulose membranes using a BioRad Trans blot turbo, 7 minutes at 25 mA. Membranes were blocked in blocking buffer (TBST with 5% milk). CDO1 was probed with anti-CDO1 rabbit polyclonal antibody (Novus # NBP2-33988), 1:500 in blocking buffer. Hif-2 was probed with anti-Hif-2 rabbit polyclonal antibody (Novus # NB100-122), 1:500 in blocking buffer. Tubulin was probed with anti-tubulin mouse monoclonal antibody (Sigma # T4026), 1:10,000 in blocking buffer. Membranes were washed 3x in TBST, 5 minutes each wash. Secondary HRP-anti-rabbit (Pierce # 31460) or HRP-anti-mouse (Pierce # 31430) were used at 1:5000 in blocking buffer. Membranes were washed 3x in TBST and developed using an enhanced chemiluminescence kit (Amersham) and exposure to film.

### NanoBiT

HEK293T cells were reverse transfected by plating 15 mL of cells at 0.625×10^6 cells/mL in complete medium (DMEM with 10% FBS) into a 75 cm^2^ flask (Corning). 90 µL of Fugene HD (BioRad #E2312) was combined with 1500 µL of Optimem I (Gibco # 31985-062) and added to 30 µg of DNA total (15 µg of HSVTK-driven VHL-LgBiT plasmid and 15 µg of CMV-driven SmBiT-CDO1 plasmid). The transfection mixture was incubated for 15 minutes at room temperature before being added drop-wise to the plated cells. After 24 hours of incubation in a cell culture incubator (37^°^C, 5% CO_2_), the transfected cells were trypsinized, normalized to 0.5×10^6 cells/mL in complete media and plated into 1536-well plates, 5 µL/well using a tip-based dispenser (GNF Systems One Tip Dispenser). The plates were incubated in a cell culture incubator (37^°^C, 5% CO_2_) for 5 hours. Next, 5 nL of 1 mM MLN4924 in DMSO was added to every well using an acoustic liquid handler (Labcyte Echo 500), and the plates were returned to the incubator for an additional 17 hours. 10 nL of compounds in 100% DMSO, or DMSO alone as a control, were dispensed into target wells using the Echo, and the plates were returned to the incubator for another 2 hours. The plates were allowed to cool to room temperature for 10 minutes, and 5 µL of NanoBiT NanoGlo reagent, consisting of NanoGlo substrate (BioRad # N2058) diluted 1:25 in a mixture of 50% NanoGlo LCS buffer (BioRad # N206C) and 50% PBS, was dispensed into each well using the GNF dispenser. After a final incubation at room temperature for 10 minutes, luminescence was read on a Viewlux plate reader (Perkin Elmer). Luminescence values were normalized using the equation (100*[well value-NC]/NC), where NC is the neutral control (DMSO). Curves were fitted using Spotfire analysis software (Tibco).

### HiBiT

HEK293T cells were reverse transfected by plating 15 mL of cells at 0.625×10^6 cells/mL in complete media (DMEM with 10% FBS) into a 75 cm^2^ flask. 90 µL of Fugene HD (BioRad #E2312) was combined with 500 µL of Optimem I (Gibco # 31985-062) and added to 30 µg of DNA total (15 µg of CMV-driven VHL plasmid and 15 µg of HSVTK-driven CDO1-HiBiT plasmid). The transfection mixture was incubated for 15 minutes at room temperature before being added drop-wise to the plated cells. After 24 hours of incubation in a cell culture incubator (37°C, 5% CO_2_), the transfected cells were trypsinized, normalized to 0.5×10^6 cells/mL in complete media and plated into 1536 plates, 5 µL/well using a tip-based dispenser (GNF Systems One Tip Dispenser). Immediately after, 5 nL of either 1 mM MLN4924 in DMSO, or DMSO alone, was added to wells using an acoustic liquid handler (Labcyte Echo 500), and the plates were incubated in a cell culture incubator (37°C, 5% CO_2_) for 5 hours. Next, 10 nL of compounds in 100% DMSO, or DMSO alone as a control, were dispensed into target wells using the Echo, and the plates were returned to the incubator for another 17 hours. The plates were allowed to cool to room temperature for 10 minutes, and 5 µL of HiBiT lytic reagent, consisting of LgBiT protein (BioRad # N4018) diluted 1:100 and lytic substrate (BioRad # N246B) diluted 1:50 in lytic buffer (BioRad # N247B), was dispensed into each well using the GNF dispenser. After a final incubation at room temperature for 10 minutes, the plates were read on a luminescence plate reader (Perkin Elmer ViewLux). Luminescence for each data point was normalized using the equation [-100*((well value-NC)/(AC-NC))], where NC is the neutral control (DMSO) and AC is the active control (mock-transfection with no DNA), and −100 corresponds to 100% degradation. Degradation AC50 values were calculated from curve fits generated by Spotfire analysis software (Tibco).

### Cell Titer Glo

The Cell-Titer Glo assay was performed identically to the HiBiT cellular protein degradation assay, except 5 µL of Cell-Titer glo substrate (BioRad # G7571) was added instead of HiBiT lytic reagent.

### Proteomics analysis of VHL molecular glue degraders: sample preparation with lysis, digestion, TMT labeling

Day 1 Huh-7 cells grown in high cysteine media (1.25 mM) were plated to 2 million cells per plate in 10 mL plated volume and incubated overnight at 37°C. Day 2, cells were treated with 5µL of 0.1 mM compounds to achieve a final concentration of 0.05 µM of compound for 24 hours. Day 3, cells were collected and washed twice in ice-cold PBS (Thermo #20012-027) and transferred as frozen cell pellets to begin proteomic analysis. Day 1 Huh-7 cells grown in high cysteine media (1.25 mM) were plated to 2 million cells per plate in 10mL plated volume and incubated overnight at 37 °C. Day 2 cells were treated with 5 µL of 0.1 mM compounds to achieve a final concentration of 0.05 µM of compound and grown in high cysteine (1.25mM) media, Day 3 cells were washed twice in ice-cold PBS (Thermo #20012-027) and transferred to our lab as frozen cell pellets to begin proteomic analysis. Cell pellets were lysed in 100 µL iST-NHS lysis buffer and sonicated to shear and break the DNA aggregates. After centrifugation, the protein concentration was measured by following a BCA™ Protein Assay Kit (Thermo 23227). All lysate protein concentrations for each condition were normalized to one concentration of 100 µg. An automated sample prep system called the PreON, developed by PreOmics, Inc. was then used to process the lysed material from digestion to TMT labeling and purification of peptides, in a fast and streamlined workflow^43,44^. The PreOn automated platform enables reproducible, high throughput sample preparation, of up to 16 samples, using reagents provided in the iST-NHS kit (PreOmics GmbH 00030). Digestion of 100 µg protein was performed for a total incubation time of 2 hrs at 37°C. Each sample was then labeled with a tandem mass tag (TMTproTM, Thermo A44522) at a ratio of 5 µg tag to 1 µg protein for a total of 90 min at room temperature. The reaction is then quenched with 0.5% hydroxylamine for 15 min. TMT-labeled samples are then combined and distributed evenly in cartridges to purify peptides. Final eluates were placed in a speed-vacuum to dry overnight.

### Liquid Chromatography Fractionation

Following elution and concentration of peptides in a Genevac EZ 2.3 Elite vacuum concentrator, approximately 530 µg of pooled multiplexed sample was fractionated by an offline high-pH fractionation system (Agilent HPLC 1200 series; Waters XBridge C18 3.5 µm, 150 mm x 2.1 mm column. Gradients are 10-40% B; Mobile phase A: 5 mM ammonium formate buffer (prepared from ammonium hydroxide), pH 10 and Mobile phase B: 90% acetonitrile, 10% of 5 mM ammonium formate buffer, pH 10). Each sample was fractionated into ninety-six fractions and then manually pooled into twenty-four final fractions. Individual fractions were subsequently concentrated and peptides reconstituted in 0.1% formic acid (FA) (to 1 µg/µL).

### Online LC-MS Chromatography

Tryptic peptides in each fraction were analyzed using an Orbitrap Fusion™ Lumos Eclipse™ Mass Spectrometer (Thermo) equipped with a IonOpticks Aurora 25 Column (1.6 µm C18, 75 µm ID x 25 cm) at an isocratic flow of 400 nL/min with a gradient of 7-28% mobile phase B (80% acetonitrile with 0.1% formic acid) in mobile phase A (0.1% formic acid) using the synchronized precursor selection (SPS) mass spectrometry to the third (MS3) mode coupled with Real Time Search (RTS) function. Briefly, the first stage of mass spectrometry (MS1) was performed in the orbitrap and scanned from 400 to 1600 mass to charge (m/z) with a resolution of 120,000. Only ions with charge state from 2+ to 6+ were selected for the second stage of mass spectrometry (MS2). The MS2 scans in the iontrap were set to a precursor isolation window of 0.7 m/z and normalized collision energy (CE) fixed at 32%. SPS-MS3 scans in the orbitrap were set to 50,000 with 10 SPS precursors, an isolation window of 2, and a CE set to 40%.

### LC-MS Data Analysis

Thermo Proteome Discoverer™ version 2.4 was used to analyze the raw mass spectrometry data. Briefly, spectra were matched against a human FASTA file downloaded from Uniprot (version July 2019 one gene/one protein with roughly 20,000 human reference proteins appended with common mass spec contaminants such as trypsin) using SequestHT algorithm. Peptide matches with <1% FDR as determined by Percolator were kept to generate a list of identified proteins. Next, accepted peptides with TMT scores as follows SPS mass matched 65%, precursor contamination 50%, minimum average reporter ion with signal/noise greater than 10 were used for protein quantitation utilizing only non-shared peptides. In-house collection of python and R scripts were used for further data normalization, e.g. adjustment for difference in total protein loaded per TMT channel, and also for assessing protein response upon compound treatment and fold change compared to DMSO treatment (n=3 biological replicates per treatment) using R Limma package (Ritchie 2015). Resulting p-values were corrected using commonly used Benjamini-Hochberg procedure. It should be noted that volcano plots show un-adjusted p-values for clarity.

### Crystallization, data collection, and x-ray structure determination

VBC complex and full length human CDO1 were dialyzed separately into a minimal buffer (5mM Tris pH 7.5, 50mM NaCl, 0.5mM TCEP) that has low ionic strength. After overnight dialysis, 1mM compound 4 or compound 8 was added to the VBC complex. After 15 minutes, hCDO1 was then added to the VBC/glue complex using 1:1 molar ratio and further incubated on ice for 15 minutes. The VBC/hCDO1/glue complex was concentrated to 20mg/ml prior to crystallization trials. A single crystallization condition was identified from a well solution containing 0.2M Sodium citrate, 0.1M Bis-tris propane pH 6.5, 20% PEG3350, after overnight incubation. Crystals were cryo-protected in reservoir solution supplemented with 20% glycerol and flash-cooled in liquid nitrogen for data collection. Datasets were collected under cryogenic conditions (100K) at the Advanced Light Source (ALS) beamline 5.0.1 and 5.0.2. All data were processed using process^45^, employing XDS^46^ or data integration and AIMLESS^47^ for scaling. Molecular replacement were carried out using PHASER^48^ with coordinates from human VHL-ElonginC-ElonginB complex (PDB ID 1VCB)^49^ and ferrous form of the cross-linked human CDO1 (PDB ID 6BGM)^50^. Structure refinement was carried out in PHENIX^51^ alternated with manual fitting in Coot^52^. Data collection and structure refinement statistics are included in Supplemental table 1. PDB IDs are VHL-4-CDO1(8VLB) and VHL-8-CDO1(8VL9).

### Protein-protein docking and WATMD analysis

All poses were simulated for 10ns in unrestrained explicit solvent at 300K, 1ATM, NTP, AM1BBC-elf charges and the PARM@FROSST force field extension on the small molecule using the AMBER suite of programs. The time averaged solvent accessible surface was determined for each pose using WATMD^20,21^. The water structure at the PPI was determined by taking the subset which was within 10 angstroms of VHL and then from that subset a further subset which is within 10 angstroms of CDO1. The resultant structure was taken as the water structure at the PPI.

## Data availability

All data generated or analyzed during this study are included in this published article (and its Supplemental files). The atomic model has been deposited in the Protein Data Bank with the accession codes 8VLB and 8VL9.

---

**Supplemental Figure 1.**
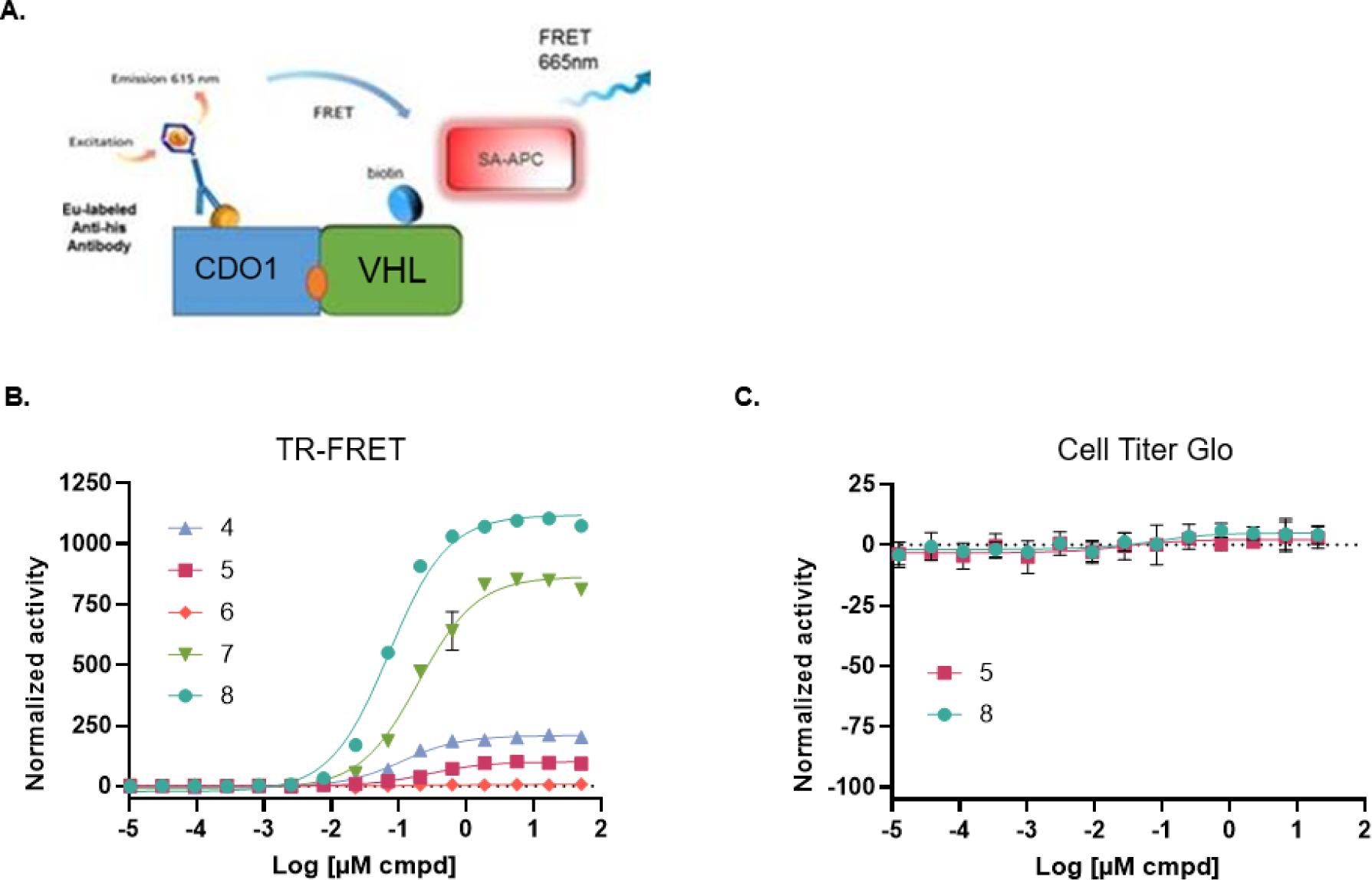
CDO1 is recruited to VHL by cpd 4 by TR-FRET. A. Schematic of TR-FRET format for biochemical VHL-CDO1 recruitment assay using anti-6His-Eu labeled donor fluorophore and streptavidin-APC acceptor fluorophore. B. TR-FRET recruitment data with compounds 4-8. Normalized activity is percent of DMSO control. C. Cell Titer Glo assay for compounds 5 and 8. Normalized activity is percent signal change relative to DMSO control.

**Supplemental Figure 2.**
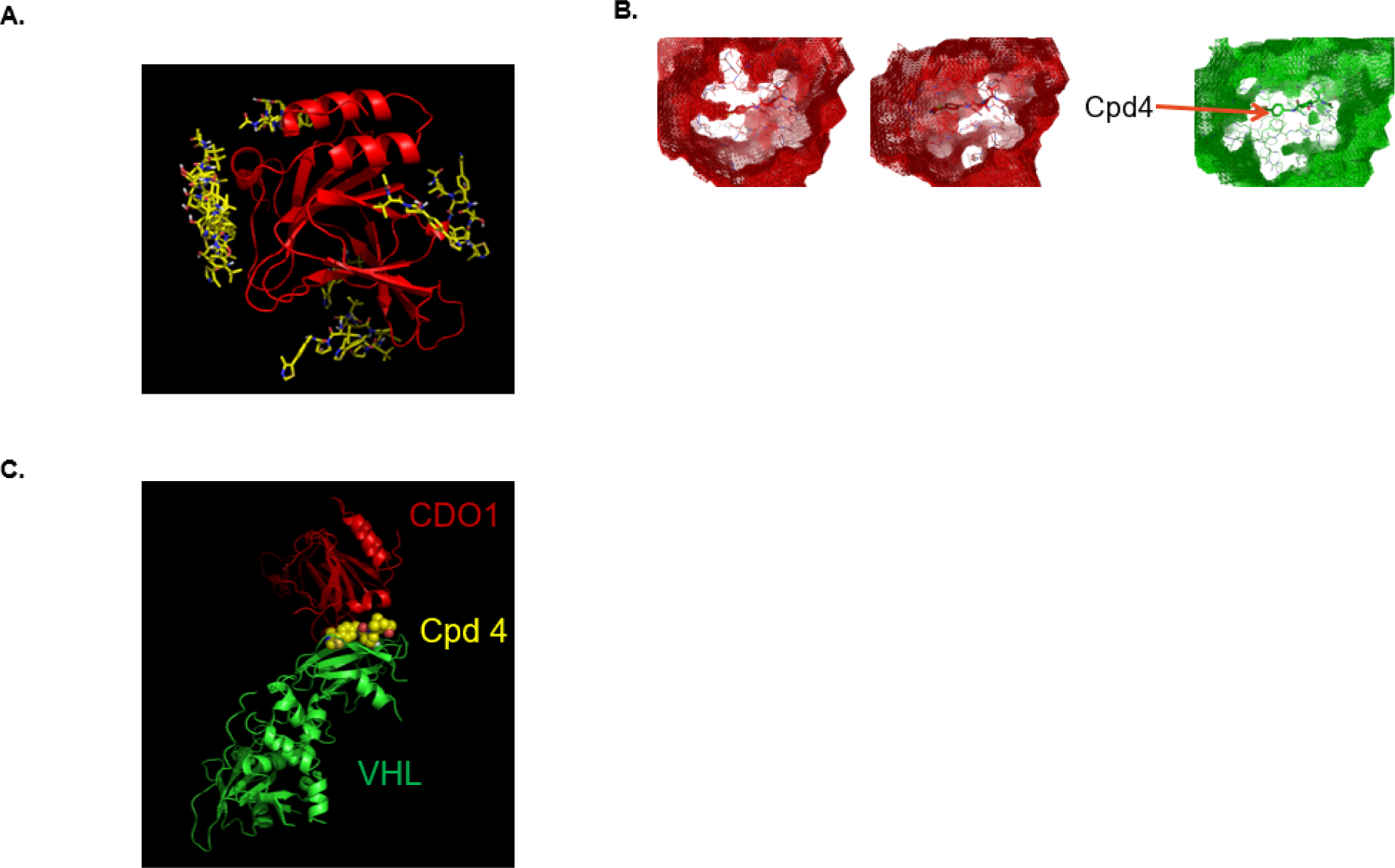
Initial prediction of CDO1-cpd interface. A. CDO1 (red) showing several clustered compound 4 (yellow) binding regions on CDO1 from docking of VHL+Cpd4 to CDO1; VHL is not shown for clarity. B. Time-averaged water-accessible surface from WATMD solvation analysis of the ternary complex in C showing the most of de-wetted region around compound 4 (right panel/green) compared to other binding modes (left panels/red) as the result of water displacement due to ternary complex formation. C. Ternary complex model from protein-protein docking with the most water displacement around compound 4 based on solvation analysis with WATMD shown in B.

**Supplemental Figure 3.**
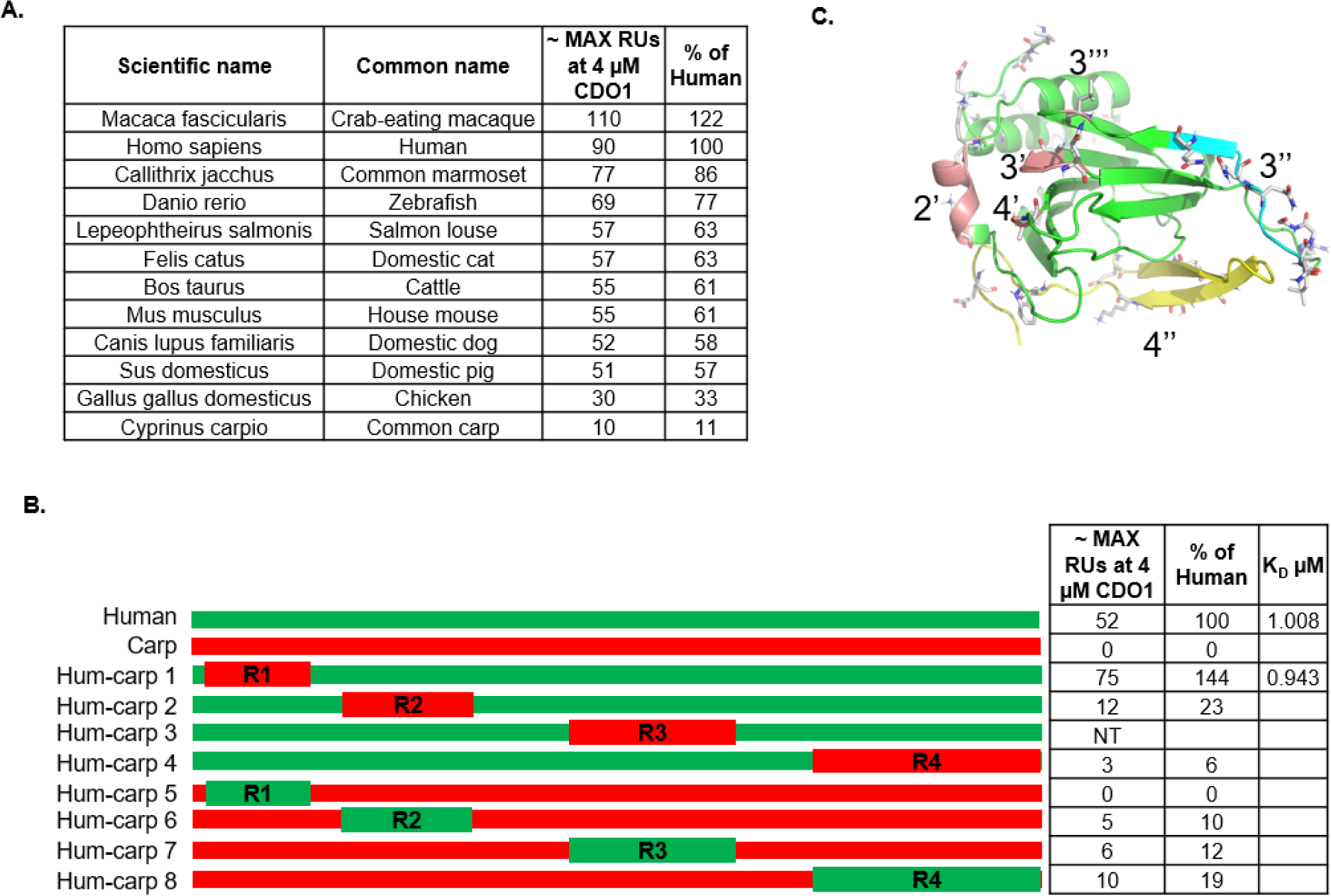
Refinement of the ternary complex interface hypothesis based on recruitment of CDO1 species orthologs and protein-protein docking analysis. A. Table of CDO1 species orthologs (n=12) chosen for this study. Indicated for each is the SPR-measured RUs (maximum observed at 4 μM CDO1) and percent activity normalized to human CDO1. B. Schematic of the first round of human-carp CDO1 chimeric proteins showing regions R1-R4 swapped between human (green) and carp (red) CDO1 proteins. Indicated for each is the SPR-measured RUs (maximum observed at 4 μM CDO1) and percent activity normalized to human CDO1. C. Refined chimera regions outlined in Fig. 3B shown on a model for potential recruitment interfaces of CDO1 based on protein-protein docking analysis and the recruitment data from chimeras outlined in B, showing refined chimeric regions 2’, 3’, 3’’, 3’’, 4’ and 4’’ as denoted and colored in Fig. 3B.

**Supplemental Figure 4.**
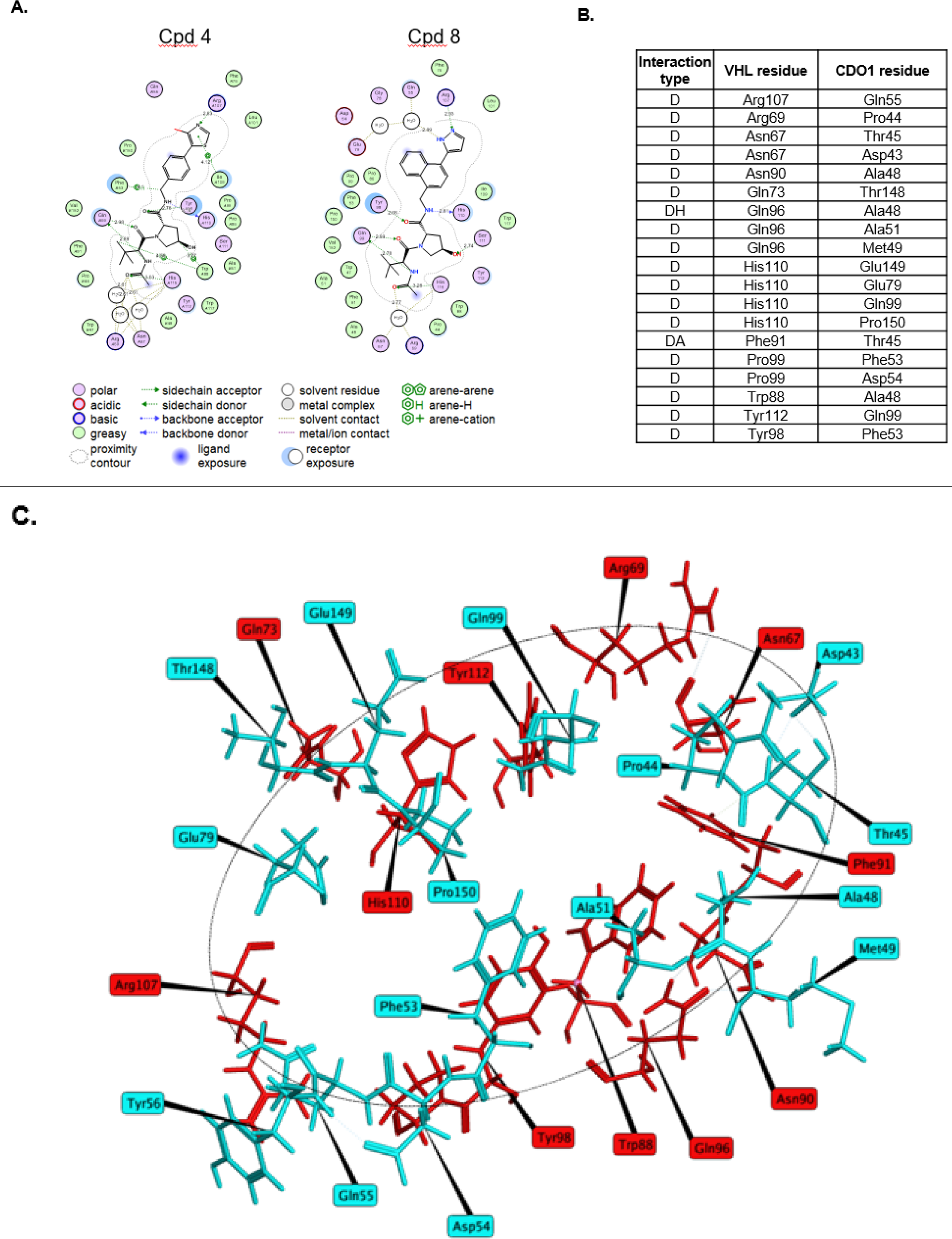
Structural details of VHL-cmpd-CDO1 ternary interactions. A. 2-D representation (MOE) of ligand interactions for both compounds 4 and 8. B. VHL-CDO1 protein-protein interactions (within 4.5 Å) shared by both the VHL-4-CDO1 and VHL-8-CDO1 ternary X-ray complexes; interaction types are distance (D), hydrogen bonding (H) and arene (A). C. Illustration showing the ring of interactions from the VHL-8-CDO1 complex. VHL residues are labeled in red and CDO1 residues are in light blue.

**Supplemental Figure 5.**
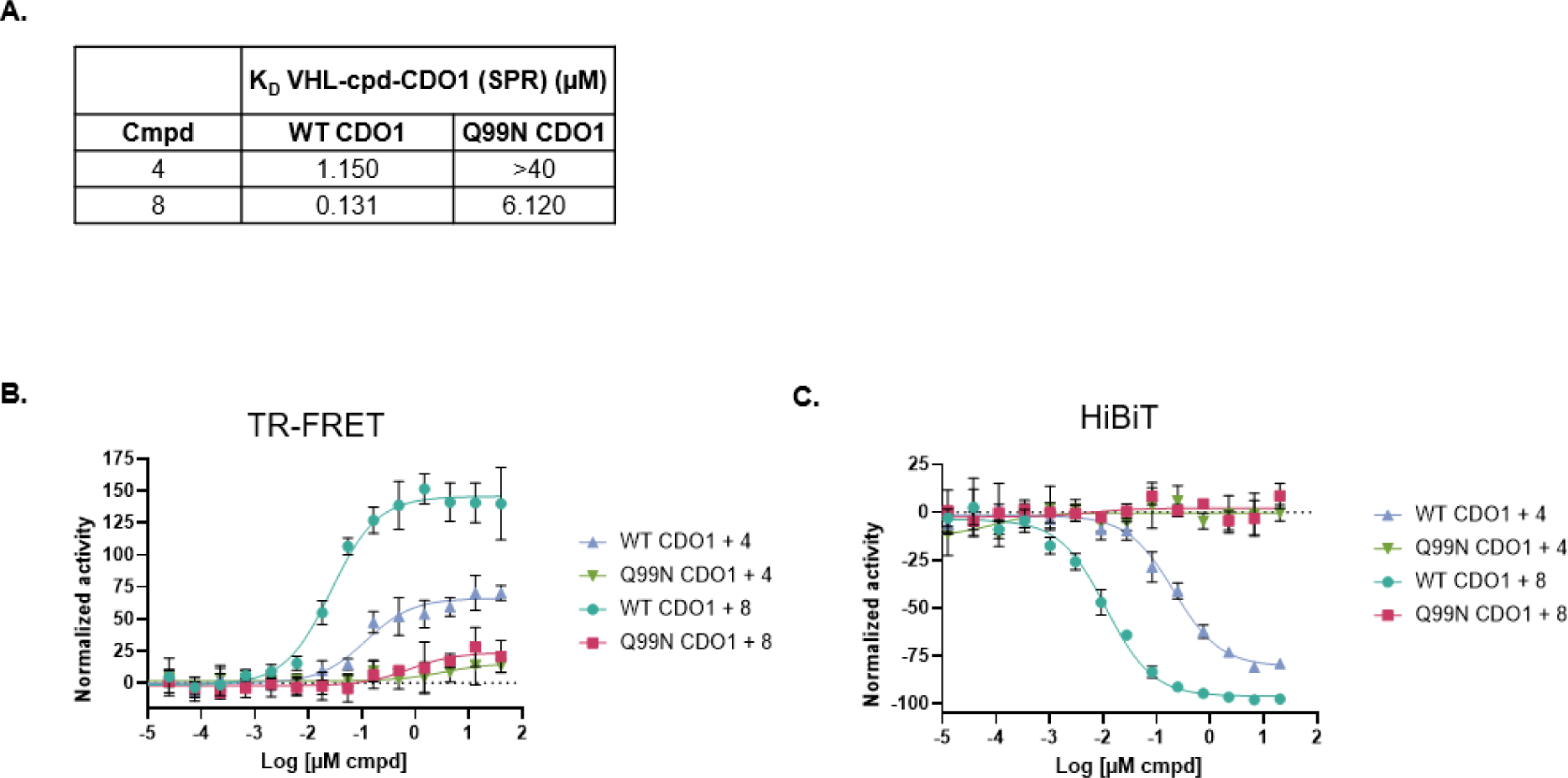
Glutamine 99 of CDO1 is critical for recruitment and degradation. A. SPR analysis of recruitment of CDO1(WT) or CDO1(Q99N) to immobilized VHL with compounds 4 or 8. K_D_ (μM) data reported from steady-state analysis. B. TR-FRET recruitment of CDO1(WT) or CDO1(Q99N) to VHL with compounds 4 or 8. Normalized activity is percent of DMSO control. C. HiBiT degradation assay for CDO1(WT) or CDO1(Q99N) with compounds 4 or 8. Normalized activity is percent signal change relative to DMSO control.

**Supplemental Figure 6.**
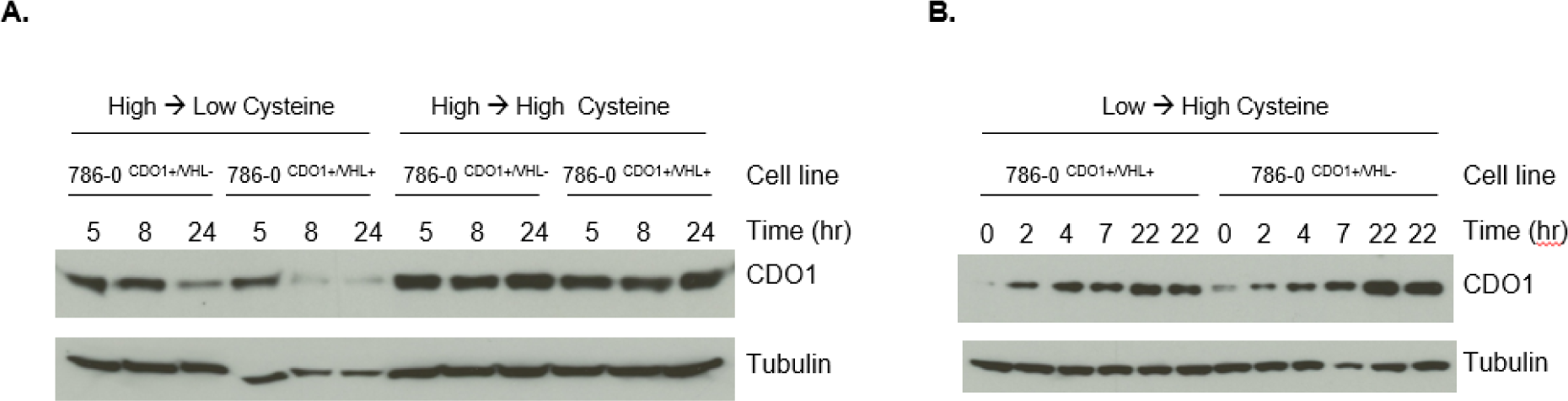
CDO1 is regulated by cellular cysteine levels independently of VHL. A. 786-0 ^CDO1+/VHL-^ (lanes 1-3, 7-9) or 786-0 ^CDO1+/VHL+^ (lanes 4-6, 10-12) cells were grown in media supplemented with high cysteine and then shifted to non-supplemented media (lanes 1-6) or maintained in high cysteine (lanes 7-12) for the indicated times before being harvested for Western blotting. B. 786-0 ^CDO1+/VHL-^ (lanes 7-12) or 786-0 ^CDO1+/VHL+^ (lanes 1-6) cells were grown in non-supplemented media and then shifted to media containing high cysteine for the indicated times before being harvested for Western blotting.

**Supplemental Figure 7.**
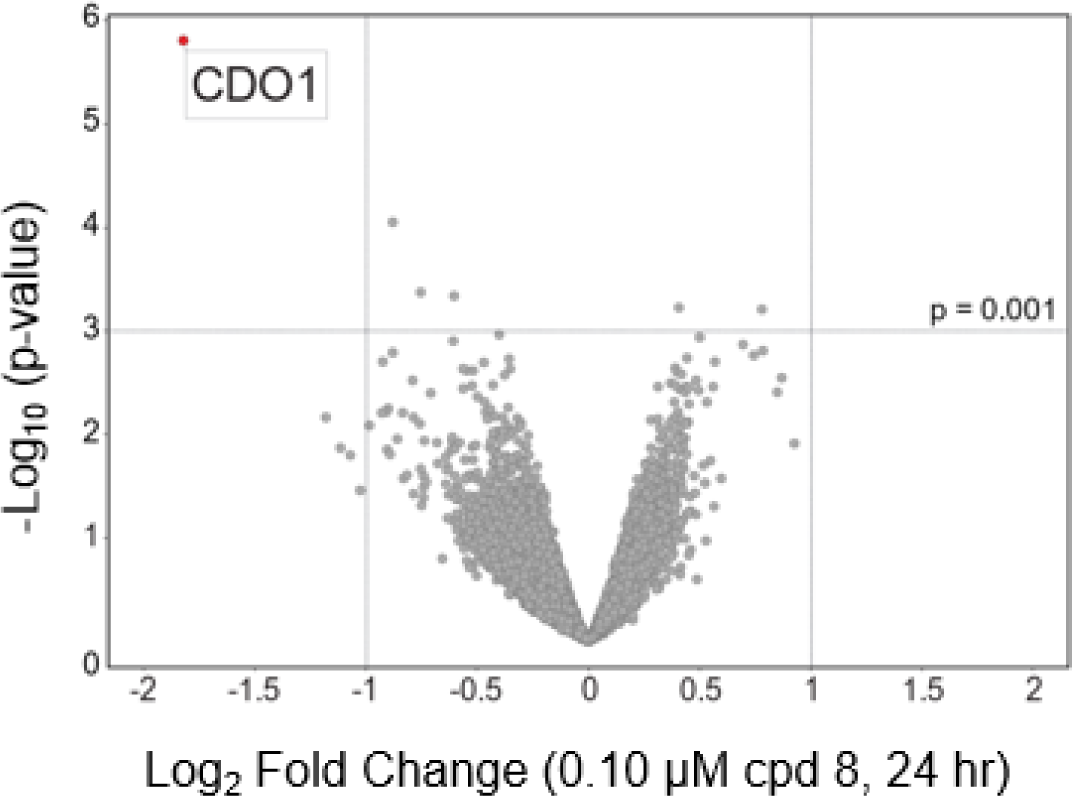
CDO1 selective downregulation at increased dose of cpd 8. Quantitative proteomics profiling of Huh-7 cells treated for 24Lhours with 0.1 µM of compound 8 or DMSO. Volcano plot shows downregulation of CDO1 upon treatment with compound 8 compared to DMSO (protein FDR < 1%, nL=L3 biological replicates per treatment). Log_2_(fold change) difference between means of treated vs. DMSO plotted against p-values calculated using Limma. Lines in the plot indicate significant cutoffs: p-value < 0.001 and absolute [Log_2_(fold change)] > 1.0. An overall degradation of 72% was observed for CDO1 at this compound treatment with biological triplicates confirming its significant modulation.

**Supplemental Table 1.**

| Supplemental Table 1 |  |  |
| --- | --- | --- |
|  | Compound 8 | Compound 4 |
| Wavelength | 1 | 1 |
| Resolution range | 38.78 - 2.5 (2.589 - 2.5) | 52.08 - 2.9 (3.004 - 2.9) |
| Space group | C 2 2 21 | C 2 2 21 |
| Unit cell | 116.657 177.873<br>103.824 90 90 90 | 116.44 178.116 104.166<br>90 90 90 |
| Total reflections | 755003 (68727) | 153748 (16552) |
| Unique reflections | 37720 (3738) | 23225 (2388) |
| Multiplicity | 20.0 (18.4) | 6.6 (6.9) |
| Completeness (%) | 99.86 (99.84) | 95.12 (99.62) |
| Mean I/sigma(I) | 20.95 (1.28) | 8.99 (0.71) |
| Wilson B-factor | 58.30 | 83.89 |
| R-merge | 0.3961 (2.324) | 0.1682 (2.58) |
| R-meas | 0.4066 (2.39) | 0.1826 (2.787) |
| R-pim | 0.09081 (0.5531) | 0.07035 (1.048) |
| CC1/2 | 0.972 (0.44) | 0.995 (0.325) |
| CC* | 0.993 (0.782) | 0.999 (0.701) |
| Reflections used in refinement | 37684 (3734) | 23188 (2379) |
| Reflections used for R-free | 1894 (190) | 1181 (129) |
| R-work | 0.1903 (0.3369) | 0.2134 (0.4236) |
| R-free | 0.2202 (0.3834) | 0.2638 (0.4361) |
| CC(work) | 0.956 (0.690) | 0.950 (0.584) |
| CC(free) | 0.924 (0.655) | 0.912 (0.544) |
| Number of non-hydrogen atoms | 4466 | 4356 |
| macromolecules | 4296 | 4304 |
| ligands | 50 | 47 |
| solvent | 120 | 5 |
| Protein residues | 533 | 533 |
| RMS(bonds) | 0.003 | 0.005 |
| RMS(angles) | 0.61 | 0.85 |
| Ramachandran favored (%) | 97.33 | 96.38 |
| Ramachandran allowed (%) | 2.67 | 3.24 |
| Ramachandran outliers (%) | 0.00 | 0.38 |
| Rotamer outliers (%) | 0.00 | 0.62 |
| Clashscore | 5.35 | 8.36 |
| Average B-factor | 70.84 | 95.32 |
| macromolecules | 71.39 | 95.61 |
| ligands | 50.64 | 73.04 |
| solvent | 59.69 | 56.05 |
| Number of TLS groups | 24 | 26 |

\*Values in parentheses are for highest-resolution shell. A single crystal was used for this structure.
\*\* $R_{\text{work}}$ and $R_{\text{free}}$ were calculated from the working and test reflection sets, respectively. The test set contained 5% of the total reflections available.

